# BAG3 Pro209 mutants associated with myopathy and neuropathy relocate chaperones of the CASA-complex to aggresomes

**DOI:** 10.1101/853804

**Authors:** Elias Adriaenssens, Barbara Tedesco, Laura Mediani, Bob Asselbergh, Valeria Crippa, Francesco Antoniani, Serena Carra, Angelo Poletti, Vincent Timmerman

**Affiliations:** Peripheral Neuropathy Research Group, Department of Biomedical Sciences, Institute Born Bunge, University of Antwerp, Antwerp, Belgium; Dipartimento di Scienze Farmacologiche e Biomolecolari, Centro di Eccellenza sulle Malattie Neurodegenerative, Università degli Studi di Milano, Milano, Italy; Department of Biomedical, Metabolic and Neural Sciences, University of Modena and Reggio Emilia, and Center for Neuroscience and Neurotechnology, Modena, Italy; VIB-UAntwerp Center for Molecular Neurology, VIB and University of Antwerp, Antwerp, Belgium

**Author notes:** These authors contributed equally (Elias Adriaenssens, Barbara Tedesco, Laura Mediani). Correspondence (V.T.), (A.P.), (S.C.).

**Keywords:** aggresome, BAG3, HSPB8, SQSTM1/p62, HDAC6, cardiomyopathy, distal myopathy, Charcot-Marie-Tooth disease type 2, chaperone-assisted selective autophagy (CASA)

## Abstract

Three missense mutations targeting the same proline 209 (Pro209) codon in the co-chaperone Bcl2-associated athanogene 3 (BAG3) have been reported to cause distal myopathy, dilated cardiomyopathy or Charcot-Marie-Tooth type 2 neuropathy. Yet, it is unclear whether distinct molecular mechanisms underlie the variable clinical spectrum of the rare patients carrying these three heterozygous Pro209 mutations in BAG3. Here, we studied all three variants and compared them to the BAG3_Glu455Lys mutant, which causes dilated cardiomyopathy. We found that all BAG3_Pro209 mutants have acquired a toxic gain-of-function, which causes these variants to accumulate in the form of insoluble HDAC6- and vimentin-positive aggresomes. The aggresomes formed by mutant BAG3 led to a relocation of other chaperones such as HSPB8 and Hsp70, which, together with BAG3, promote the so-called chaperone-assisted selective autophagy (CASA). As a consequence of their increased aggregation-proneness, mutant BAG3 trapped ubiquitinylated client proteins at the aggresome, preventing their efficient clearance. Combined, these data show that all BAG3_Pro209 mutants, irrespective of their different clinical phenotypes, are characterized by a gain-of-function that contributes to the gradual loss of protein homeostasis.

## Introduction

Protein homeostasis is maintained by a complex network of molecular chaperones and co-chaperones providing protection to client proteins at every stage of their life-time (Balchin et al. 2016). As soon as a nascent polypeptide leaves the ribosomal exit tunnel, chaperones interact with exposed domains to facilitate protein folding (Gloge et al. 2014). In case of protein misfolding, chaperones will either try to refold or guide the polypeptide towards degradation by proteasomes or the autophagy-lysosomal pathway (Balchin et al. 2016).

The activity of many chaperones is critically dependent on co-chaperones. One family of co-chaperones is represented by the Bcl2-associated athanogene (BAG) family of proteins, which in humans include six members, encoded by 6 different genes (Takayama and Reed, 2001). All six family members share a conserved BAG-domain, which is essential for their binding to the Hsp70 chaperones (Behl, 2016). BAG3 is a well-characterized family member that contains a number of additional protein domains besides the conserved BAG-domain, including two Ile-Pro-Val (IPV)-motifs, a PxxP domain and a WW-domain (**Fig. 1a**). Each of these domains is known to have specific interacting partners. For instance, the BAG-domain is known to mediate the interaction with Hsp70/Hsc70 or Bcl2 (Takayama et al. 1995, 1997 and 1999). The IPV-motifs have been shown to be indispensable for binding to small heat shock proteins (sHSPs) (Fuchs et al. 2010), the WW-domain binds LATS1 (Meriin et al. 2018), and the PxxP domain is necessary for the interaction with dynein and PLC-γ (Doong et al. 2000, Gamerdinger et al. 2011).

**Fig. 1.**
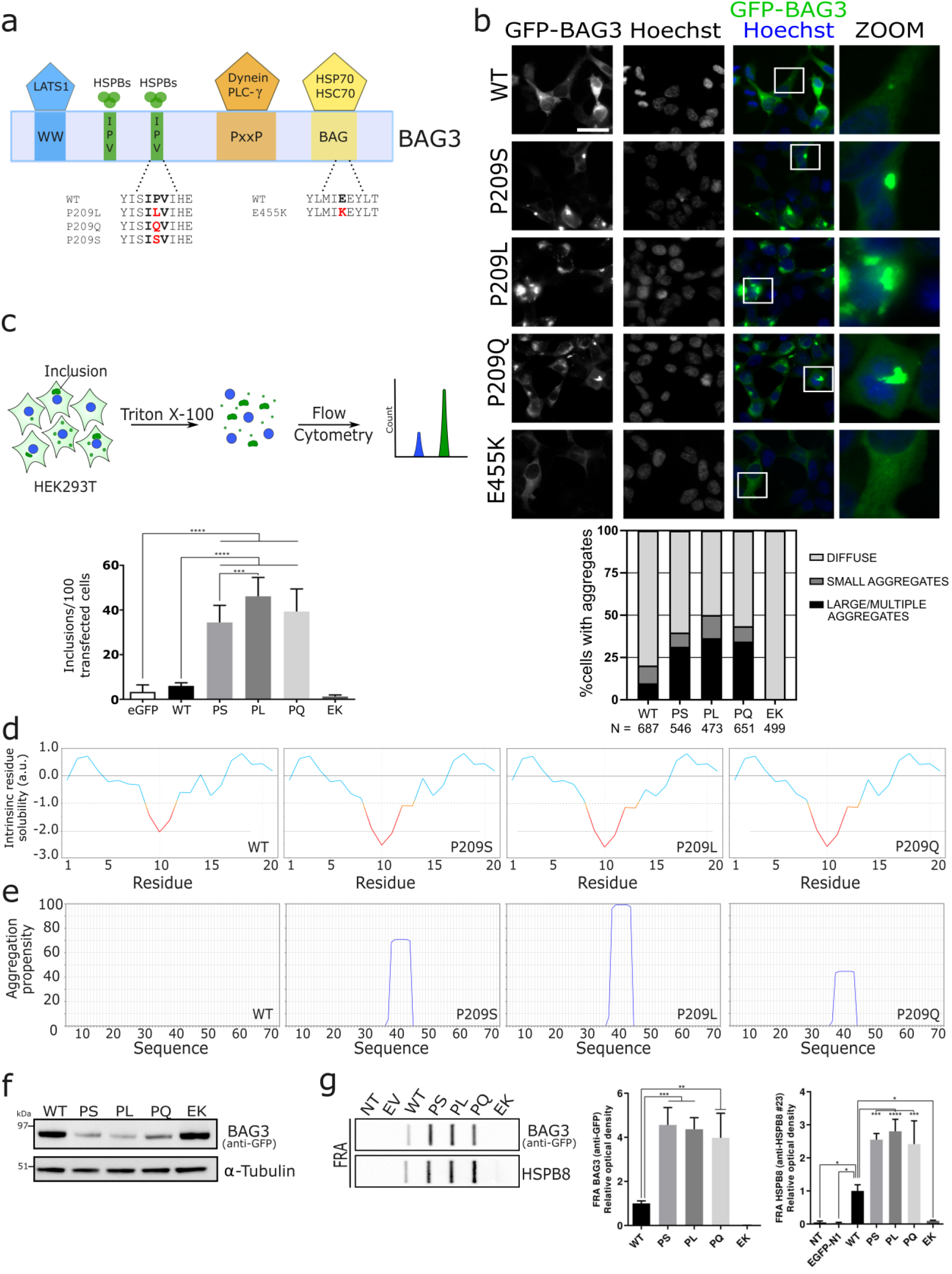
BAG3_Pro209 mutations cause cytoplasmic aggregation. (**a**) Schematic representation of the structure of BAG3, including the WW-domain, the two IPV-motifs, the PxxP-domain and BAG-domain. The known interactors of each motif are shown at the top and the missense mutations that were studied in this manuscript are shown at the bottom in red. (**b**) HEK293T cells stably expressing HSPB8-V5 were transiently transfected with BAG3-GFP constructs. Six random fields were selected for analysis. The mean number of cells counted per field was 95 and thus over 400 cells per genotype were counted. (scale bar = 10 μm) (**c**) Quantification of BAG3-GFP inclusions using Flow cytometric analysis of inclusions (FloIT). Transiently transfected HEK293T cells were collected and stained with DAPI prior to 0.1% Triton X-100 treatment. The intracellular BAG3-GFP inclusions and Hoechst-positive nuclei are subsequently quantified using flow cytometry. Bar graph represents the means of BAG3-GFP cytoplasmic inclusions per 100 transfected cells. One-Way ANOVA with Bonferroni’s multiple comparisons test were used for statistical analysis. (**d-e**) Bio-informatic analysis of (**d**) the solubility of wild type or mutant BAG3 with CamSol and (**e**) of the aggregation propensity with Tango software. (**f**) Western blot analysis of the NP-40 soluble fraction from HEK293T cells stably expressing HSPB8-V5 and transiently transfected with BAG3-GFP constructs. The constructs were abbreviated as followed: wild type (WT), Pro209Ser (PS), Pro209Leu (PL), Pro209Gln (PQ), Glu455Lys (EK). One of three representative western blots is shown. (**g**) Filter-retardation assay (FRA) analysis of the NP-40 insoluble fraction. Anti-GFP and anti-HSPB8 antibodies were used to detect insoluble levels of BAG3 (wild type or mutants) and HSpB8. Relative optical densities are reported in the graphs as means ± SD of normalized values. One-Way ANOVA with Bonferroni’s multiple comparisons test were used for statistical analysis (n=3). The constructs were abbreviated as followed: non-transfected (NT), empty vector (EV), wild type (WT), Pro209Ser (PS), Pro209Leu (PL), Pro209Gln (PQ), Glu455Lys (EK).

The sHSP with the highest affinity for the IPV-motifs of BAG3 is HSPB8 (Hsp22) (Morelli et al. 2017, Rauch et al. 2017). In fact, the protein stability of HSPB8 is critically dependent on BAG3, as it is rapidly degraded in its absence (Carra et al. 2008). As also other members of the HSPB family are capable of binding to BAG3, it is thought that in case HSPB8 would be unable to fulfil its role (e.g. due to lower expression levels of HSPB8), these other sHSPs could partly replace its function by binding to BAG3. Such compensatory mechanisms would ensure that BAG3-sHSP interactions are maintained even under compromising conditions and underscore the importance of this interaction.

BAG3-HSPB8 is a sub-complex at the basis of a larger protein complex, known as the chaperone-assisted-selective-autophagy (CASA) complex. In addition to BAG3 and HSPB8, also Hsp70/Hsc70, CHIP and SQSTM1/p62 are part of this complex (Arndt et al. 2010). Under stress conditions, unfolded or misfolded proteins are rapidly recognized by sHSPs (Carra et al. 2019), which are then transferred to ATP-dependent chaperones like Hsp70 for refolding. In case the substrate cannot be refolded by Hsp70, then the client is directed towards autophagosomal degradation by CHIP (an E3 ubiquitin ligase that ubiquitinates the substrate (Jiang et al. 2001, Murata et al. 2001, McDonough et al. 2003)) and SQSTM1/p62 (an autophagy receptor that binds to ubiquitinated proteins and transfers them to autophagosomes (Lamark et al. 2005, Ciuaffa et al. 2015, Cha-Molstad et al. 2017)). By clustering the different components into a single complex, substrates are likely handed over faster to reduce the potentially dangerous dwell time.

The complete substrate repertoire of the CASA-complex remains elusive. However, many model client proteins involved in the appearance of adult onset neurodegenerative diseases were shown to aggregate less in the presence of the CASA-complex, such as elongated polyQ-variants of the mutant Androgen receptor (AR), mutant Huntingtin (HTT) or mutant ataxin-3 (ATX-3), SOD1 mutants, TDP43 mutants, neurotoxic dipeptides deriving from the expanded repeat of the C9ORF72 mRNA. At the very least this illustrates that the CASA chaperone complex can handle a diverse array of misfolded proteins (Rusmini et al. 2017).

In light of its important role for the maintenance of cellular protein homeostasis, it is not surprising that mutations in the co-chaperone *BAG3* have been reported to cause a variety of disorders affecting distal muscles, cardiomyocytes or peripheral nerves. One hot-spot residue is the proline at codon 209 of BAG3. Genetic variants of this codon were previously linked to cardiomyopathy and distal myopathy (Selcen et al. 2009, Semmler et al. 2014). More recently, also two families with late-onset axonal Charcot-Marie-Tooth (CMT) neuropathy were reported with a novel Pro209Ser mutation in *BAG3* (Shy et al. 2018). The impact of the different Pro209 mutations on the function of BAG3 remains unclear. The Pro209Leu mutant, which causes early-onset dilated cardiomyopathy and/or severe distal myopathy, was shown to cause a toxic gain-of-function by impairing the Hsp70 client processing (Meister-Broekema et al. 2018). This results in the accumulation of aggregated proteins that sequester important protein quality control (PQC) factors, including Hsp70. Importantly, accumulation of aggregated proteins has been documented in the biopsies of patients affected by myopathy and peripheral neuropathy (Selcen et al. 2009, Weihl et al 2018), further supporting the interpretation that these mutations may affect BAG3 PQC functions.

Here, we studied the impact of the three heterozygous Pro209 missense mutations and compared it with the cardiomyopathy-causing BAG3_Glu455Lys mutant, which is located in the BAG-domain (Villard et al. 2011), and the wild type BAG3 protein. We found that all BAG3_Pro209 mutants equally interact with the other CASA-components compared to wild-type. However, due to their increased propensity to aggregate, all BAG3_Pro209 mutants acquire a toxic gain-of-function leading to the sequestration of all CASA components, along with their bound polyubiquitinated clients, in perinuclear aggregates called aggresomes. This, in turn, promotes a general collapse in the chaperone-network, as previously reported for the P209L mutant (Meister-Broekema et al. 2018).

## Results

We studied the pathogenic consequences of 3 previously reported BAG3 missense mutations, located in the second IPV-motif of BAG3 (**Fig. 1a**) which mediates the interaction with sHSP family members, and causing distinct phenotypes: Pro209Leu (early-onset dilated cardiomyopathy and/or severe distal myopathy (Selcen et al. 2009)), Pro209Gln (late-onset distal myopathy (Semmler et al. 2014)), Pro209Ser (late-onset CMT2 (Shy et al. 2018)). In addition, we included the clinical and molecular well-characterised BAG3_Glu455Lys mutant (Fang et al. 2017). The BAG3_Glu455Lys mutation is located in a different protein-domain of BAG3 (**Fig. 1a**), the BAG-domain which mediates the direct interaction with Hsp70, and causes dilated cardiomyopathy (Villard et al. 2011).

### BAG3 Pro209-mutations cause BAG3 protein aggregation

To investigate the impact of the three IPV-mutations, we transiently overexpressed GFP-tagged wild type and mutated BAG3 in HEK293T cells that stably overexpress HSPB8. The expression of exogenous wild type BAG3 compared to endogenous BAG3 is about 5-fold higher after transient transfection of these HEK293T cells (**Fig. S1**). Of note, the HSPB8-BAG3-Hsp70 complex identified in HeLa cells has a stoichiometry corresponding to 2:1:1 (Carra et al., 2008). Thus, to maintain this stoichiometry, we stably overexpressed HSPB8 in HEK293T cells, which are characterized by low expression levels of HSPB8 and abundant Hsp70.

We performed fluorescence microscopy using these HEK293T cells that stably overexpress HSPB8 to verify the subcellular distribution of BAG3. The images showed a different localization and distribution of the three IPV-motif located BAG3_Pro209 mutants compared to wild type BAG3 and the BAG-domain located BAG3_Glu455Lys mutant (**Fig. 1b**). Both Glu455Lys and wild type BAG3 showed a predominantly diffuse cytoplasmic distribution (**Fig. 1b**). In contrast, BAG3_Pro209 mutants showed an aberrant distribution, with low levels of diffuse cytoplasmic BAG3 protein and higher levels of BAG3-GFP protein sequestered in multiple smaller aggregates or one irregular shaped large perinuclear aggregate. More specifically, 31.4% of cells transfected with Pro209Ser mutant, 36.5% of cells transfected with Pro209Leu and 34.4% of cells transfected with Pro209Gln mutant presented with large aggregates at 24 hours after transfection (**Fig. 1b**). From this quantification, subtle differences were detected between the three Pro209 mutants; as the Pro209Leu mutation caused aggregation in a slightly higher number of cells compared to the other mutants.

To confirm these results with an independent technique, we made use of a recently developed method to quantify cellular protein aggregates in a high-throughput manner, known as FloIT (Whiten et al. 2016). This method employs the fluorescence counting capabilities of a flow cytometer to determine the number of cellular aggregates in cellular lysates (**Fig. 1c**). HEK293T cells that stably overexpress HSPB8 were transiently transfected with the different BAG3-GFP proteins, lysed in a mild detergent (0.1% Triton X-100 in PBS) supplemented with DAPI for nuclear staining. GFP-positive aggregates were then investigated by FloIT (**Fig. S2**). All three IPV-mutants formed significantly more inclusions than wild type BAG3 or the BAG-domain Glu455Lys mutant (**Fig. 1c**). Similar to what we observed with fluorescence microscopy, the Pro209Leu mutant formed a higher amount of aggregates. Together, these data demonstrate that all three mutants affecting the IPV-motif cause protein aggregation, a phenotype that seems unique to IPV-mutants, as the BAG-domain Glu455Lys mutant and BAG3 wild type protein remained diffusely distributed in the cytoplasm.

To assess if this altered cytoplasmic distribution also affects the solubility of the protein, we first employed two bio-informatic prediction tools: CamSol (Sormanni et al. 2015) and Tango (Fernandez-Escamilla et al. 2004), to determine whether the IPV-mutants reduce protein solubility. Both methods predicted a strong reduction in protein solubility for each of the genetic mutants **(Fig. 1d-e and Fig. S3**). Interestingly, both prediction methods suggested that the Pro209Leu substitution would reduce protein solubility the most. To determine whether these mutants are indeed less soluble, we extracted the proteins from cells overexpressing wild type or mutated BAG3 using a buffer that contains 0.5% of NP-40 as detergent (**Fig. 1f**). While the protein levels of the Glu455Lys mutant were very similar to those of wild type BAG3 in detergent soluble fractions, the levels of all three BAG3_Pro209 mutations (Ser, Leu and Gln) were drastically decreased in the NP-40 buffer. To confirm that these mutants become insoluble, we used a filter-retardation assay (FRA). This showed that the Pro209 mutants were highly enriched in the NP-40 insoluble fraction (**Fig. 1g**). Interestingly, also HSPB8 was found in higher amounts in the insoluble fraction. Summation of both soluble and insoluble fractions shows that total levels of BAG3 are slightly increased for mutants (**Fig. S4**).

Since the mutations affect the heart, muscle or peripheral motor neurons in patients, we verified whether the phenotypes described above are also observed in these cell types. To this end, we overexpressed GFP-tagged BAG3 in mouse myoblasts (C2C12 cells) and immortalized motor neurons (NSC-34 cells). The results obtained in these two cell lines were identical to the ones obtained in HEK293T cells: the BAG3_Pro209 mutants also aggregated in C2C12 and NSC-34 cells, while BAG3 wild type or BAG3_Glu455Lys did not (**Fig. 2**).

**Fig. 2.**
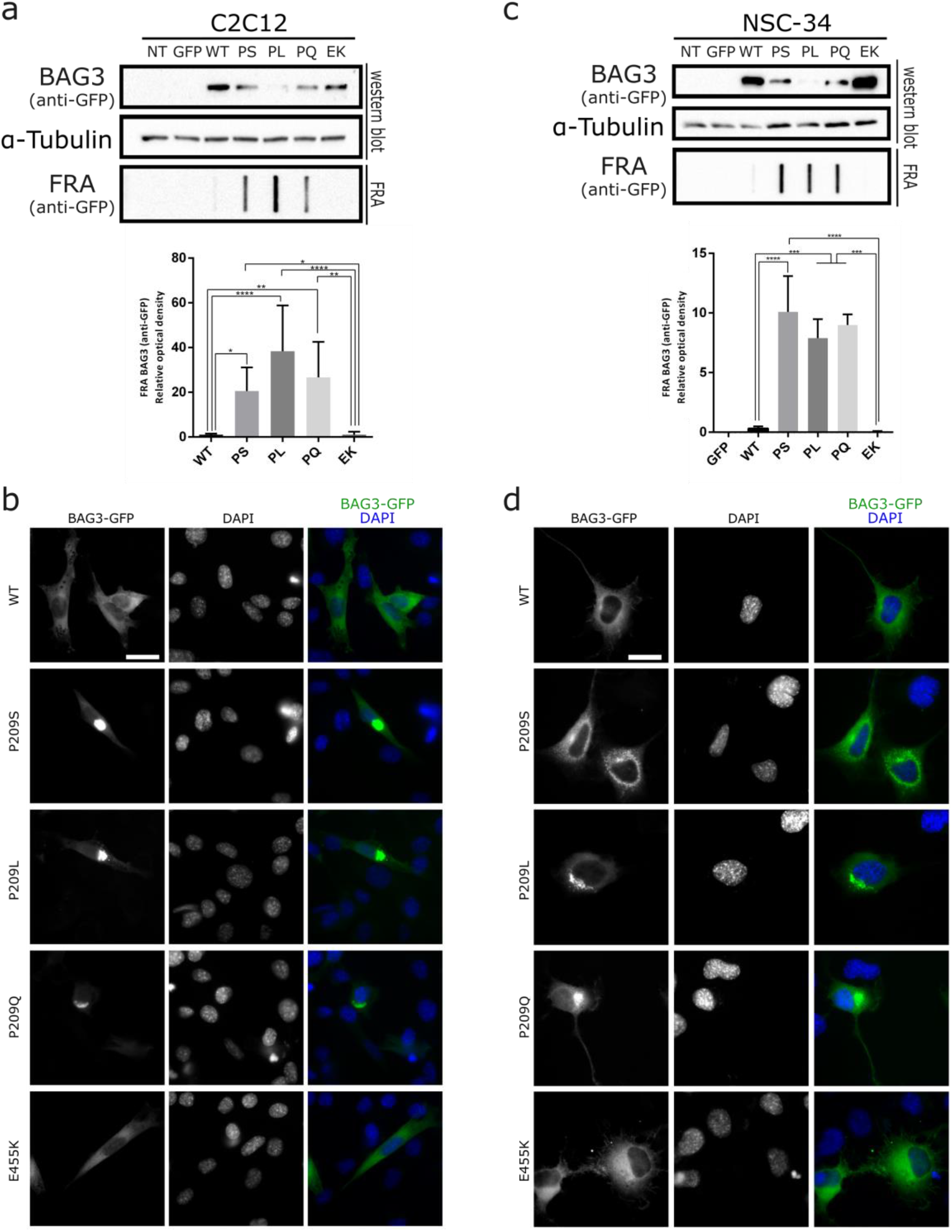
BAG3_Pro209 mutants also aggregate in muscle (C2C12) and motoneuron-like cells (NSC-34). We transiently transfected GFP-tagged BAG3 wild type or mutant constructs in C2C12 and NSC-34 cells. We then verified protein aggregation by separating the soluble fraction (western blot) and insoluble fraction (filter retardation assay (FRA)) (**a** and **c**) or verified protein aggregation by immunofluorescence (**b** and **d**). The FRA analysis is displayed for the NP-40 insoluble fraction. Relative optical densities are reported in the graphs as means ± SD of normalized values. One-Way ANOVA with Bonferroni’s multiple comparisons test were used for statistical analysis (n=3). Scale bar = 10 μm.

Combined these data demonstrate that all BAG3_Pro209 mutants have a decreased protein solubility and this gives rise to large protein aggregates in the cytosol, regardless of the cell type investigated.

### The perinuclear aggregates formed by BAG3 Pro209 mutants are aggresomes

Since BAG3 aggregates have an irregular shape and BAG3 was previously shown to translocate to aggresomes (Gamerdinger et al. 2011), we assessed whether these structures were aggresomes. As HDAC6 and vimentin are two well-known markers for aggresomes (Johnston et al. 1998, Kawaguchi et al. 2003), we verified whether the BAG3_Pro209 variants colocalized with HDAC6 or vimentin. Confocal images confirmed that the BAG3_Pro209 aggregates are positive for both HDAC6 (**Fig. 3a**) and vimentin (**Fig. 3b**), supporting the interpretation that BAG3_Pro209 mutants accumulate in the form of aggresomes.

**Fig. 3.**
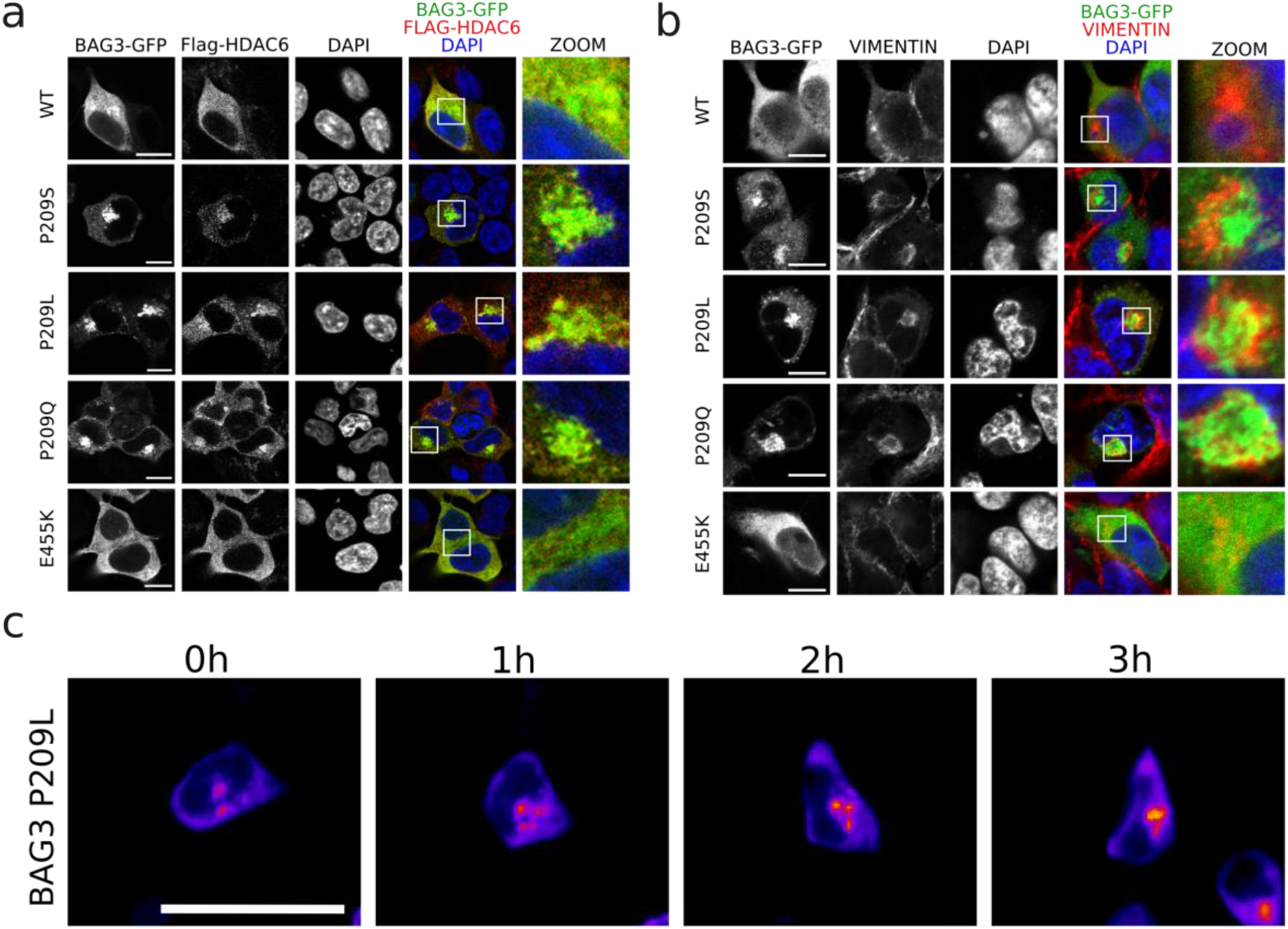
BAG3_Pro209 mutants accumulate at aggresomes. Co-localization was assessed between BAG3-GFP and aggresome-markers in HEK293T cells stably expressing HSPB8-V5 and transiently transfected for 24 h with BAG3-GFP constructs. As markers for aggresomes we used: (**a**) FLAG-HDAC6, and (**b**) endogenous vimentin. Scale bar = 10 μm. (**c**) Live-cell time-lapse imaging of GFP-tagged BAG3-Pro209Leu in HEK293T cells. Scale bar = 25 μm.

To gain insight into the different stages of this process, we performed live-cell time-lapse imaging in HEK293T cells after transient transfection of GFP-tagged BAG3_Pro209Leu. We observed that mutant BAG3 first formed smaller aggregates at the periphery of the cell; with time these smaller aggregates clustered in one central spot near the nucleus (**Fig. 3c**). Similar results were obtained in HeLa cells, further suggesting that aggregation is an intrinsic property of this mutant (**Fig. S5**).

### BAG3 Pro209 mutants sequester chaperones of the CASA-complex at aggresomes

BAG3 forms a stoichiometric complex with HSPB8 and Hsp70/Hsc70 and the Pro209 residue is located within the binding domain of HSPB8 to BAG3 (Carra et al. 2008, Fuchs et al. 2010). We therefore verified by co-immunoprecipitation whether the Pro209 mutations affect the ability of BAG3 to bind its binding partners HSPB8 and Hsp70/Hsc70 (**Fig. 4a and Fig. S6**). Our data show that none of the BAG3 mutants abolished the interaction with HSPB8, nor with Hsp70/Hsc70 (**Fig. 4a and Fig. S6**). The interaction with Hsp70/Hsc70 was only affected by the BAG3_Glu455Lys mutation, which is located within the BAG domain essential for binding Hsp70 (Fang et al. 2017, Rauch et al. 2016). By performing the reverse experiment, assessing the co-immunoprecipitation of BAG3 along with HSPB8, we obtained similar results (**Supplementary Fig. S7**). By contrast, the interaction with the CASA-partner SQSTM1/p62 was increased by the Pro209 mutations compared to wild type BAG3 (**Fig. 4a**). This is consistent with a previous independent report (Guilbert et al. 2018). So, although the IPV-motifs mediate the interaction with HSPB8, we found that Pro209 mutants primarily affect the interaction with SQSTM1/p62, which, as far as we know, is not mediated by a direct interaction between the two proteins, but requires the assembly of the full CASA-complex bound to poly-ubiquitin chain linked misfolded proteins.

**Fig. 4.**
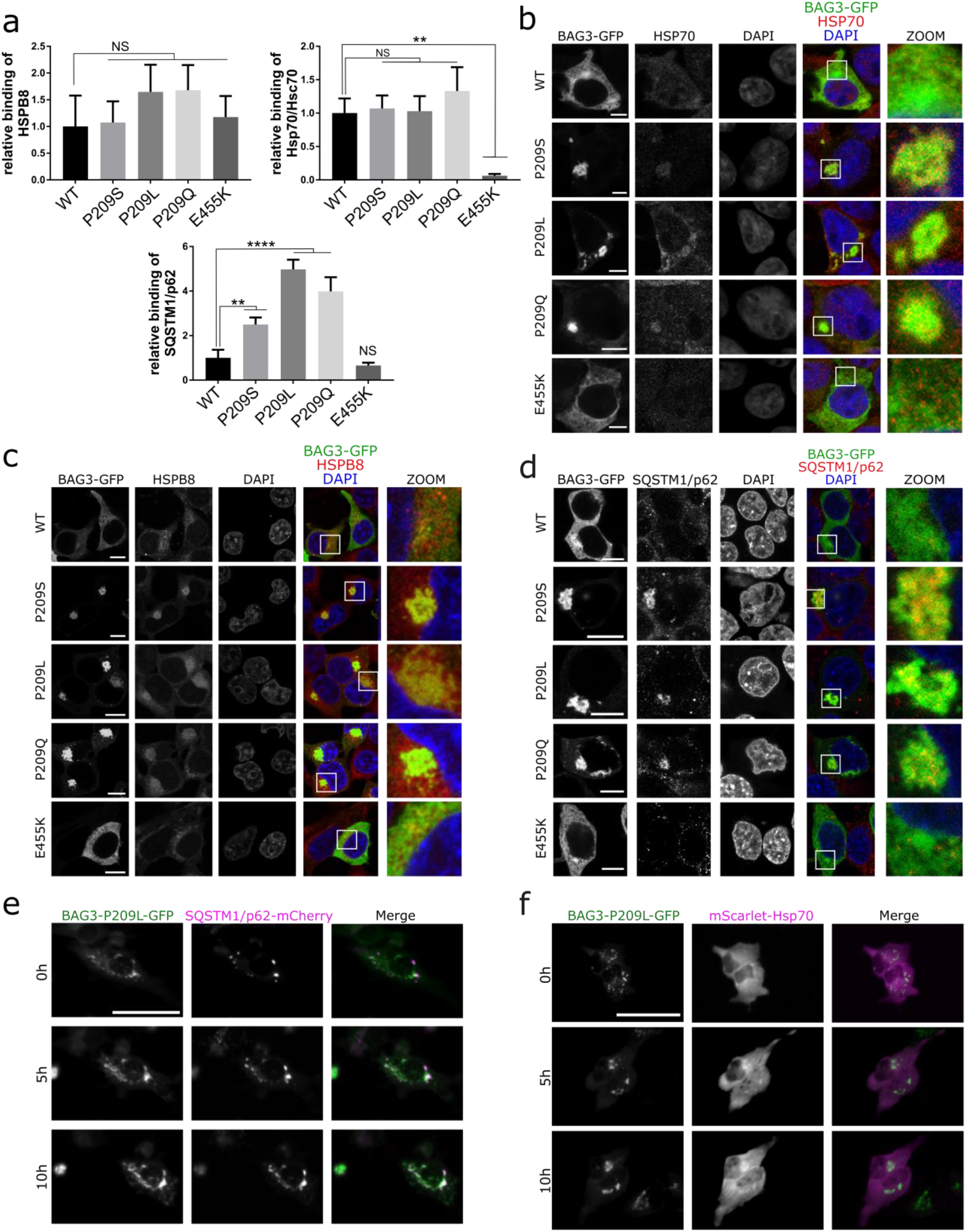
BAG3_Pro209 mutants sequester other members of the CASA-complex in aggresomes. HEK293T cells that stably overexpress HSPB8-V5 were transiently transfected with wild type or mutant BAG3-GFP constructs to assess the interaction between BAG3 and components of the CASA-complex. (**a**) Co-immunoprecipitation of BAG3-GFP and the CASA-complex using the GFP-trap system. The amount of interacting proteins was quantified and corrected for the amount of immunoprecipitated BAG3 as represented in the graph bar (means ± SD). One-Way ANOVA with Bonferroni’s multiple comparisons test were used for statistical analysis. NS = non-significant, ** p<0.01, and **** p<0.0001 (n=3). The wild type (WT) or mutants were abbreviated as followed: Pro209Ser (PS), Pro209Leu (PL), Pro209Gln (PQ), Glu455Lys (EK). (**b-d**) Immunocytochemistry of BAG3-GFP constructs to assess colocalization with (**b**) endogenous Hsp70, (**c**) HSPB8, and (**d**) SQSTM1/p62. Scale bar = 5 μm (b) and 10 μm (c and d). (**e-f**) Live-cell time-lapse imaging of GFP-tagged BAG3-Pro209Leu and RFP-tagged SQSTM1/p62 or Hsp70. HeLa cells were transiently transfected with mutant BAG3-GFP constructs and (**e**) mCherry-tagged SQSTM1/p62 or (**f**) mScarlet-tagged Hsp70. Cells were imaged once per hour. Scale bar = 50 μm.

To assess whether BAG3 relocates the CASA-complex to aggresomes, we performed co-localization experiments. Hsp70 and HSPB8 showed a diffuse cytoplasmic distribution in cells expressing BAG3 wild type or the Glu455Lys mutant (**Fig. 4b-c**). By contrast, in cells overexpressing BAG3_Pro209 mutants, we observed relocalization of both Hsp70 and HSPB8 to aggresomes (**Fig. 4b-c**), in line with the transition of HSPB8 from the NP-40 soluble to the insoluble fraction in a manner similar to BAG3 (**Fig. 1g**).

Next, we tested if mutant BAG3 aggresomes were also positive for sequestosome 1 (SQSTM1/p62). SQSTM1/p62 has the ability to bind both ubiquitin and protein degradation machinery and was found to regulate the formation of aggresomes (Lamark et al. 2005, Fujita et al. 2011, Ciuaffa et al. 2015, Cha-Molstad et al. 2017). Using confocal imaging, we observed that SQSTM1/p62 indeed colocalizes with the perinuclear BAG3 aggregates formed by all three Pro209 mutants, while SQSTM1/p62 maintained its typical disperse pattern in cells overexpressing wild type or Glu455Lys mutant BAG3 (**Fig. 4d**). So other members of the CASA-complex relocate to the aggresome in cells expressing BAG3_Pro209 mutants.

To investigate when Hsp70 and SQSTM1/p62 are recruited to the BAG3-aggregates, we performed live-cell time-lapse imaging of transiently transfected HeLa cells. We co-transfected GFP-tagged BAG3 with mCherry-tagged SQSTM1/p62 or mScarlet-tagged Hsp70. We found that both Hsp70 and SQSTM1/p62 co-localized with BAG3 in pre-aggresome bodies, which were transported over time towards the maturing aggresome (**Fig. 4e-f**); this supports the interpretation that Hsp70 and SQSTM1/p62 associate with BAG3 already in the early stages of the aggregation process. Note that we could not verify HSPB8, as tagging the small protein with a fluorescent protein of the same size, could potentially interfere with its functioning.

In summary, as a consequence of its increased aggregation propensity, mutant BAG3_Pro209 relocates Hsp70, HSPB8 and SQSTM1/p62 to the aggresomes, potentially decreasing their availability and compromising their functioning.

### BAG3 Pro209 mutants are trapped at aggresomes due to slower subunit exchange between the soluble and insoluble fraction

BAG3 wild type was shown to be degraded mainly by autophagy (Arndt et al. 2010). We confirm that both BAG3 wild-type and the Pro209Leu mutant are degraded through autophagy (**Fig. S8**). Thus, the increased aggregation propensity of the BAG3_Pro209Leu mutants and their accumulation at the aggresome, may compromise their turnover. To this end, we performed a cycloheximide wash-out experiment, which allowed us to determine the degradation rate of BAG3. We transfected HEK293T cells, that stably overexpress HSPB8, with wild type or mutant BAG3-GFP constructs, and subjected those cells to cycloheximide treatment for different time points. At time point zero, 6 and 12 hours after cycloheximide treatment, we collected protein lysates and separated the soluble from non-soluble fractions to be able to specifically study the protein turnover of the non-soluble aggresome-enriched fraction. First, in line with **Fig. 1**, we found that BAG3_Pro209 mutants were enriched in the non-soluble fraction (**Fig. 5a**). However, wild type BAG3 had a comparable depletion curve in the soluble and insoluble fraction (**Fig. 5a and Fig. S9**). In contrast, mutant BAG3 was depleted faster in the soluble fraction than in the insoluble fraction suggesting either increased protein degradation of mutant BAG3 or increased transitioning of mutant BAG3 from soluble to insoluble.

**Fig. 5.**
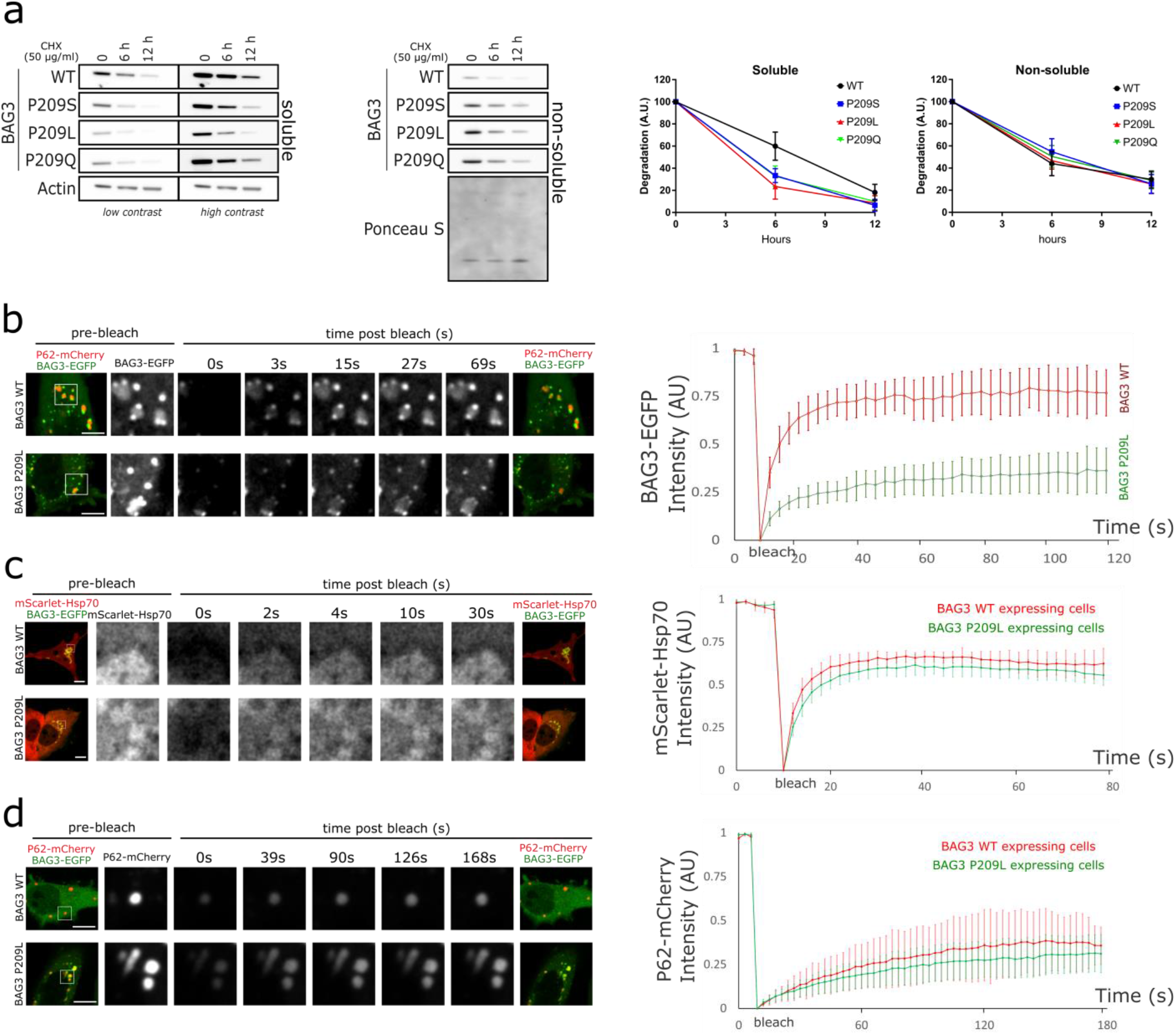
BAG3_Pro209 mutants are trapped in long-lasting aggresome structures due to reduced subunit exchange. (**a**) Protein degradation rates were determined with a cycloheximide wash-out experiment. HEK293T cells that stably overexpress HSPB8-V5 were transiently transfected with wild type or mutant BAG3-GFP constructs and subjected to cycloheximide treatment (50μg/ml) for the indicated time. Protein turnover of BAG3-GFP was determined by western blot after separation of the soluble from insoluble fraction. (n=3). (**b-d**) Fluorescence recovery after photobleaching (FRAP) analysis was performed on HeLa cells that were transiently transfected with BAG3-GFP and mScarlet-Hsp70 or SQSTM1/p62-mCherry constructs. Bleaching was performed either on (**b**) BAG3-GFP, (**c**) mScarlet-Hsp70, or (**d**) SQSTM1/p62-mCherry. Quantification of the fluorescence intensity over time was plotted for cells overexpressing WT and mutant BAG3. Graph bar shows the means (± SD) over time (n=6). Scale bar = 10 μm

To gain further insights into the dynamics of BAG3 at aggresomes, we performed fluorescence recovery after photo-bleaching (FRAP) experiments on HeLa cells overexpressing wild type or mutant BAG3-GFP. This allowed us to assess if aggresomes still exchange BAG3-subunits with the pool of soluble cytosolic proteins, providing information on the solubility of these inclusions. Note that in cells overexpressing wild type BAG3, the number of cells with aggresome-like structures is very low, as we showed in **Fig. 1**. However, for the FRAP experiments, we specifically selected this minority of cells in order to be able to compare the recovery rates in BAG3-positive inclusions. The FRAP-measurements demonstrated that wild type BAG3 recovered rapidly after photobleaching of small cytoplasmic inclusions, demonstrating its dynamic behaviour and rapid exchange between compartments (**Fig. 5b**). For mutant BAG3, we observed that a large proportion of pre-aggresomal BAG3_Pro209Leu mutant is immobile. Furthermore, compared to wild type BAG3, the exchange rate of the mobile BAG3_Pro209Leu mutant subunits is drastically slower (**Fig. 5b**). We noted that there seem to be two types of pre-aggresome bodies: one type is positive for SQSTM1/p62 and the other type is negative for SQSTM1/p62. We therefore repeated the FRAP experiment and compared the recovery rate of BAG3_Pro209Leu in SQSTM1/p62-positive pre-aggresomes versus SQSTM1/p62-negative pre-aggresomes. The fluorescence recovery of BAG3 was not different between SQSTM1/p62-positive and SQSTM1/p62-negative pre-aggresome bodies (**Fig. S10**), indicating that the presence of SQSTM/p62 is not influencing BAG3 mobility.

As our data show that mutant BAG3 is trapped in aggresomes and that our co-localization data show that other members of the CASA-complex are also present in these aggresomes, we verified whether mutant BAG3 also disturbed the subunit exchange rate of other members of the CASA-complex. We therefore performed FRAP on aggresome-like structures in cells overexpressing wild type or mutant BAG3 and found that neither Hsp70 nor SQSTM1/p62 had altered fluorescence recovery rates, which suggests that their subunit exchange and mobility is not altered by mutant BAG3 (**Fig. 5c-d**).

Together, these data show that two distinct pools of mutant BAG3 exist: one pool of mutant BAG3 is trapped in aggresome-associated structures with drastically reduced subunit exchange compared to wild type BAG3, while a second pool of mutant BAG3_Pro209Leu is moving freely within the cytosol. Due to a reduced exchange with the cytosolic (soluble) fraction, initial engagement with pre-aggresome bodies commits mutant BAG3 towards the aggresome, where it holds a residence time in the range of hours. This process occurs independently of SQSTM1/p62 recruitment at the BAG3 pre-aggresome bodies.

### BAG3 Pro209 mutants reduce the chaperone-capacity of the CASA-complex

Meister-Broekema et al. (2019) showed that BAG3_Pro209 mutants fail to stimulate Hsp70-dependent client processing, leading to the sequestration of ubiquitinylated Hsp70-bound clients into aggregates. We verified whether the aggresomes formed by all BAG3_Pro209 mutants were enriched for ubiquitinylated proteins, which would suggest a failure to degrade Hsp70-bound clients.

To this end, we transiently transfected HEK293T cells that stably overexpress HSPB8-V5 and separated the soluble from the insoluble fraction. We found that the insoluble fraction from cells expressing BAG3_Pro209 mutants contained more ubiquitinylated proteins (**Fig. 6a**). Interestingly, the amount of ubiquitinylated proteins in the soluble fraction was the same as for wild type BAG3. Using confocal microscopy, we confirmed that these insoluble ubiquitinylated proteins cluster at the aggresome (**Fig. 6b**). This suggests that clients are still recognized by the CASA-complex, but that a failure in client processing leads to accumulation of ubiquitinylated proteins at aggresomes. This failure has been suggested to have important implications for cell function and disease. For example, the CASA complex facilitates the removal of filamin, which is essential for muscle maintenance (Arndt et al. 2010). Importantly, the BAG3_Pro209Leu mutant is unable to properly clear damaged filamin; this, in turn leads to its accumulation in form of aggregates, contributing to muscle cell dysfunction in BAG3_Pro209Leu patients (Arndt et al. 2010).

**Fig. 6.**
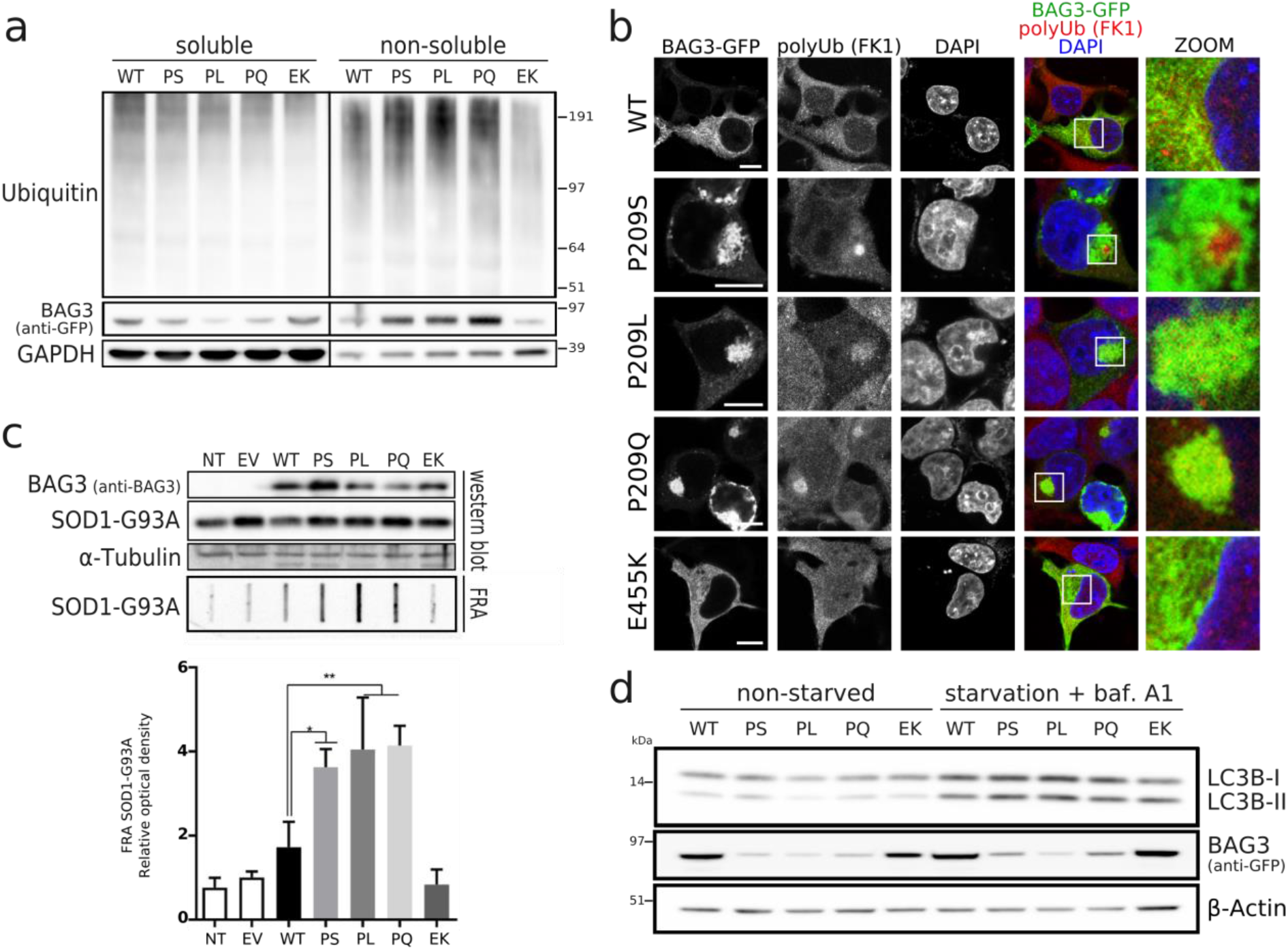
BAG3_Pro209 mutations cause a failure in chaperone-function of the CASA-complex. Chaperone-activity was assessed in HEK293T cells stably expressing HSPB8-V5 and transiently transfected with wild type or mutant BAG3-GFP constructs. (**a**) Aggregation of ubiquitinated clients was verified by separation of the soluble and insoluble fraction. Both fractions were analysed by western blot with anti-ubiquitin as marker for the accumulation of ubiquitinylated-proteins. (**b**) Immunocytochemistry of BAG3-GFP constructs to assess colocalization with ubiquitinated proteins. Scale bar = 10 μm (**c**) Protein aggregation assay by transient transfection of model client protein SOD1-G93A. The same total protein lysates were analysed by western blot and filter-retardation assay (FRA). Relative optical densities are reported in the graph as means ± SD of normalized values. One-Way ANOVA with Bonferroni’s multiple comparisons test were used for statistical analysis. * p<0.05, ** p<0.01 (n=3). (**d**) Autophagic activity was determined by western blot, before and after starvation by serum depletion plus 10nM bafilomycin A1 for 2 hours. Protein lysates were analysed by SDS-PAGE with LC3B-II as a marker for autophagosomes. Following abbreviations were used: non-transfected (NT), empty vector (EV), wild type (WT), and BAG3 mutations Pro209Ser (PS), Pro209Leu (PL), Pro209Gln (PQ), Glu455Lys (EK).

To further test the hypothesis that the BAG3_Pro209 mutants acquire a toxic gain of function that ultimately impairs the clearance of aggregation-prone proteins, we studied the degradation of a well-characterized model client known to be targeted for autophagy-mediated clearance by the CASA-complex (Crippa et al 2010). To this end, we co-transfected SOD 1-G93A together with wild type or mutant BAG3. While the soluble levels of SOD1-G93A were similar in cells expressing the different BAG3 variants, we detected a significantly higher amount of SOD1-G93A in the insoluble fraction in cells expressing the three Pro209 mutants (**Fig. 6c**). We next tested another known CASA-complex substrate, the peptide poly-GA, an aggregation-prone dipeptide repeat protein produced from the ALS-linked C9orf72 gene (Cristofani et al. 2017). Similar to SOD1-G93A, the degradation of poly-GA was impaired in cells overexpressing BAG3_Pro209 mutants (**Fig. S11**).

So far our data argue against the possibility that failure to degrade their clients by BAG_Pro209 mutants is due to the inability of the CASA-complex to recognize them (ubiquitinylated proteins accumulate at the aggresome, which suggests that the clients are still correctly recognized by the CASA-complex). Alternatively, the client is recognized and bound by the BAG3_Pro209 mutants, but it is no longer released for degradation by the autophagosomes; another possibility would be that the BAG3_Pro209 mutants impair the autophagy degradation pathway. We first verified whether the autophagic flux was impaired, since the aggresome is highly enriched in autophagosomal structures and this route is used for client degradation. As shown in **Fig. 6d**, the autophagic pathway was not impaired by BAG3_Pro209 mutations, suggesting that the accumulation of ubiquitinylated proteins cannot be explained by impairment in autophagy and supporting the idea that the CASA-complexes composed of BAG3_Pro209 mutants fail to release the bound client from Hsp70 for degradation by autophagosomes. This interpretation is in line with Meister-Broekema et al. (2019), who showed that BAG3_Pro209Leu fail to stimulate Hsp70-dependent client processing.

### HDAC6 interference does not prevent aggresome formation by BAG3 Pro209 mutants

Since the BAG3_Pro209 mutations lead to accumulation of ubiquitinylated clients at aggresomes due to a failure in client degradation, we verified whether interference with the aggresome-formation pathway could be pursuit as a therapeutic strategy. To this end, we focused on the histone deacetylase HDAC6 for two reasons: (i) as it was previously shown that HDAC6 is essential for aggresome formation upon proteasome inhibition (Kawaguchi et al. 2003), and (ii) since HDAC6-inhibitors have shown promising results as a therapeutic strategy in the field of motoneuron and neuromuscular disorders (d’Ydewalle et al. 2011, Guo et al. 2017, Benoy et al. 2018, Helleputte et al. 2018, Prior et al. 2018).

We inhibited HDAC6 with Tubastatin A, which is an inhibitor that binds HDAC6 specifically but has no activity towards other HDACs (d’Ydewalle et al. 2011), and we verified the aggresome formation and protein aggregation in HEK293T cells stably overexpressing HSPB8-V5 and transiently transfected with wild type or mutant BAG3 constructs. To ensure HDAC6 was inhibited prior to the aggresome formation by mutant BAG3, we started the Tubastatin A treatment two hours before transfection of wild type or mutant BAG3 plasmids. The effectiveness of the treatment was confirmed by the increase in tubulin acetylation, as HDAC6 is well known to deacetylate tubulin (Hubbert et al. 2002, Matsuyama et al. 2002). However, HDAC6 inhibition with Tubastatin A did not prevent the aggregation of mutant BAG3, the accumulation of ubiquitinylated proteins in the insoluble fraction, or the formation of aggresomes (**Fig. 7a-b**).

**Fig. 7.**
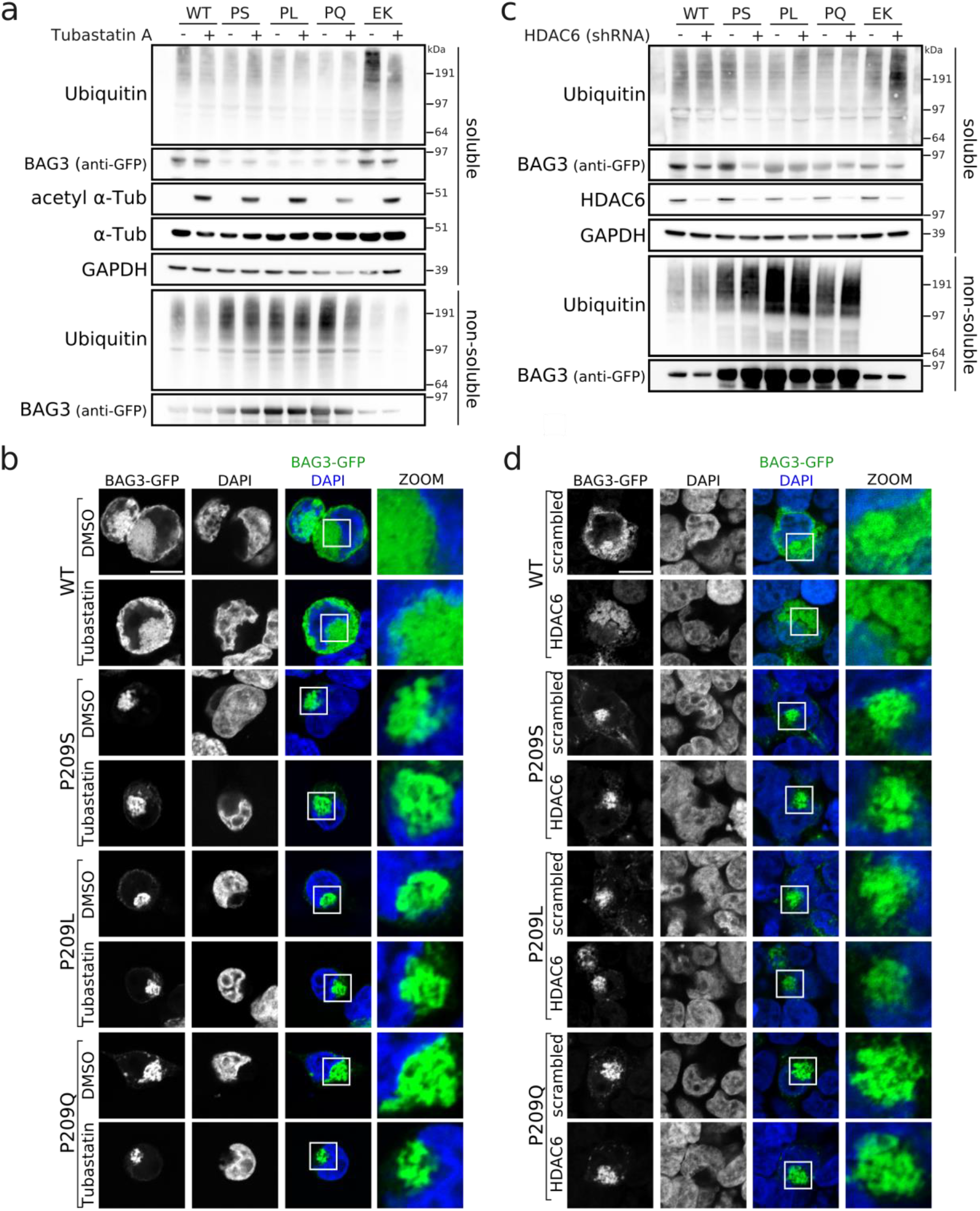
HDAC6-inhition with tubastatin A or HDAC6-depletion with shRNA does not rescue BAG3_Pro209-associated phenotypes. The protein aggregation and aggresome formation of BAG3-Pro209 mutants was assessed in HEK293T cells stably expressing HSPB8-V5 and transiently transfected with wild type or mutant BAG3-GFP constructs before and after HDAC6 inhibition (**a-b**), or depletion by shRNAs (**c-d**). Following abbreviations were used: wild type (WT), and BAG3 mutations Pro209Ser (PS), Pro209Leu (PL), Pro209Gln (PQ), Glu455Lys (EK). Scale bar = 10 μm

Although Tubastatin A effectively inhibited the deacetylase function of HDAC6, we wanted to rule out that other protein domains of HDAC6 were still contributing to aggresome formation. To this end we generated a stable knockdown line for HDAC6 by lentivirally transducing a short hairpin RNA in HEK293T cells stably expressing HSPB8-V5. We expressed wild type or mutant BAG3 in this HDAC6-knockdown line and, despite the drastic reduction in HDAC6 protein levels, the protein aggregation of mutant BAG3, the accumulation of ubiquitinylated client proteins in the insoluble fraction, and the formation of aggresomes were not prevented by depletion of HDAC6 (**Fig. 7c-d**).

Therefore, neither pharmacological inhibition nor genetic depletion of HDAC6 prevented aggresome formation in BAG3_Pro209 mutant cells. Inhibition of HDAC6 may therefore not offer the desired therapeutic potential to rescue the compromised chaperone-function in cells expressing BAG3_Pro209 mutants. Moreover, these data suggest that BAG3_Pro209 mutants induce aggresome formation downstream of HDAC6 or from an independent pathway.

## Discussion

Aggresome formation is a cellular response to an overload of misfolded proteins (Johnston et al. 1998). It involves many components from PQC factors, such as SQSTM1/p62 and chaperones, to cytoskeletal elements such as γ-tubulin and vimentin. The latter seem required for the clustering of the misfolded proteins (Johnston et al. 1998). This effort to group misfolded proteins at one well-determined spot ensures that potentially toxic proteins are removed from the remaining cytosol and protects the cell from adverse effects. The aggresome is therefore rich in ubiquitinylated proteins and requires chaperones and autophagosomes to remove and degrade these components in a controlled manner.

BAG3 is a scaffolding constituent that clusters different components of the PQC system into one protein complex. Upon inhibition of proteasomes, the BAG3-complex becomes activated and translocates to aggresomes to deliver ubiquitinylated proteins for degradation (Gamerdinger et al. 2011). In this study, we found that disease-associated BAG3-mutations of Pro209 decrease the protein solubility leading to the aggregation of BAG3 and associated factors (**Fig. 8**). As a consequence, this leads to the formation of aggresomes which are not only rich in BAG3 but also Hsp70, HSPB8 and ubiquitinylated substrates. We found that a reduction in exchange of BAG3 between soluble and insoluble pre-aggresomal puncta underlies the clustering of BAG3 at the aggresome. Due to this slower exchange, the initial engagement of BAG3 with non-soluble compartments commits BAG3 towards the forming aggresome. As such, BAG3 but also Hsp70, HSPB8 and the ubiquitinylated substrates that are bound by Hsp70 and HSPB8 are all transported towards the aggresome. This leads to clustering of ubiquitinylated species at the aggresome where a failure in the Hsp70-cycle, due to mutations in BAG3 as shown by Meister-Broekema et al. (2019), prevents the ubiquitinylated proteins from being degraded.

**Fig. 8.**
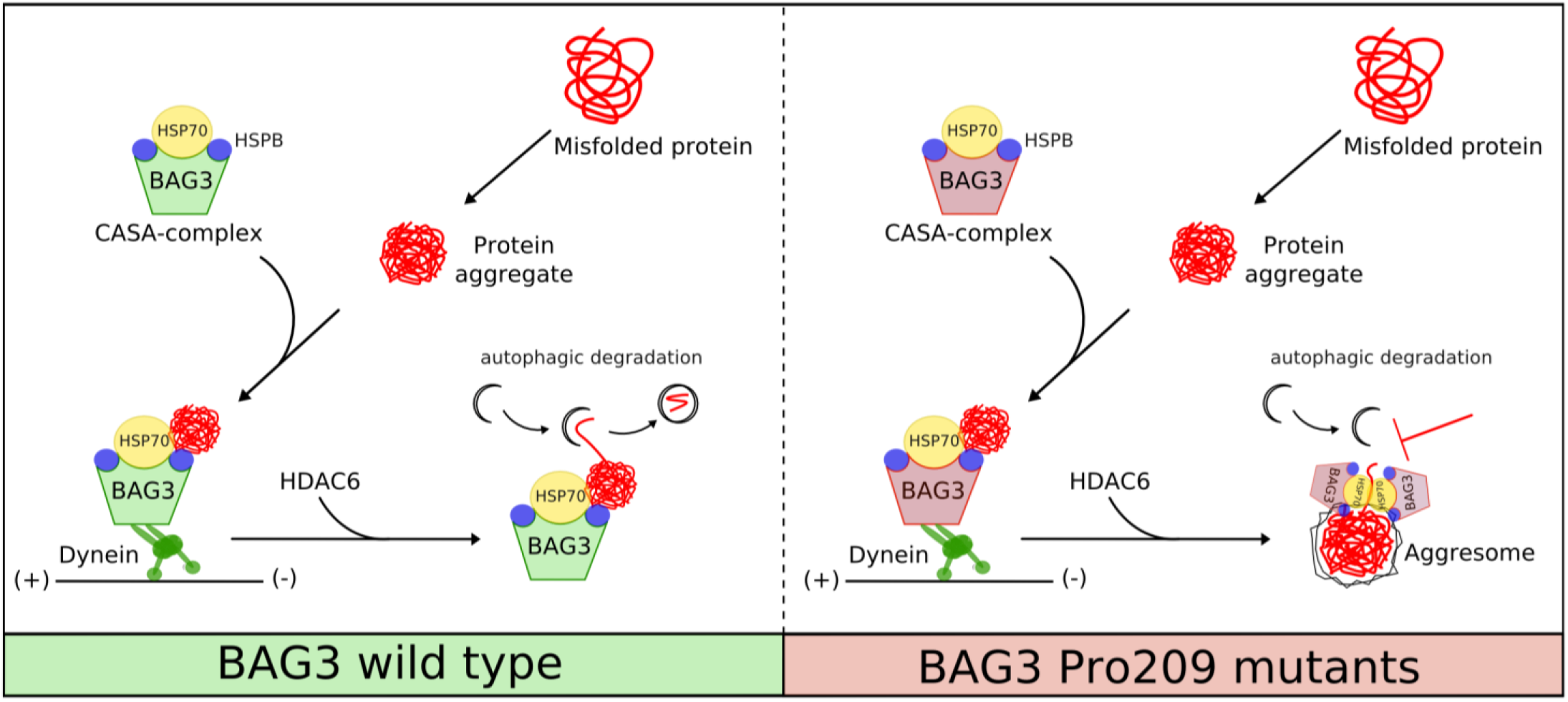
Schematic summary. Misfolded proteins are captured by the CASA-complex and transported to the MTOC, where autophagosomes are concentrated and efficiently degrade the misfolded cargo. BAG3_Pro209 mutations destabilize the protein’s intrinsic stability and lead to BAG3 aggregation. Moreover, due to a toxic gain-of-function, Pro209Leu BAG3 impairs the functional chaperone-cycle of Hsp70 (Meister-Broekema et al., 2018). As a consequence, the CASA-complexes that contain mutant BAG3 accumulate at the aggresome with its bound clients and co-factors, preventing on the one hand the degradation of the Hsp70-bound misfolded cargo and sequestering important proteostasis factors such as HSPB8, SQSTM1/p62 and ubiquitin.

The chaperone-failure of the Hsp70 processing cycle is a surprising finding given that the mutations reside in a highly conserved motif for sHSP binding, while not affecting the HSPB8-BAG3 association (Meister-Broekema et al. 2019). Of note, also binding of BAG3 to Hsp70 is not affected by the Pro209 mutations (this study and Meister-Broekema et al. 2019). This raises two important questions. (i) Is the processing of HSPB8-specific clients also affected by the BAG3_Pro209 mutations? The fact that mutations in the *HSPB8* gene are linked to muscle atrophy, together with the finding that the function and stability of HSPB8 depend on BAG3 (Carra et al. 2008), may suggest that altered Hsp70-BAG3 mediated processing of HSPB8-specific clients may have an impact on skeletal muscle function. (ii) To which extent do the IPV-motifs contribute to the chaperone-function of the CASA-complex? One way to test this would be by developing a mouse model that has the two IPV-motifs in BAG3 deleted, similarly to what has been developed for *in vitro* experiments (Fuchs et al. 2010). This may then provide new insights in the diverse compositions and functions of the CASA-complex and help in understanding why IPV-mutations give rise to such diverse clinical phenotypes.

Noteworthy is how similar the different Pro209 mutants behave in our cellular and biochemical assays, despite their distinct clinical phenotypes. The only difference was that the Pro209Leu has a mildly increased propensity to aggregate compared to the two other Pro209 mutants; the Pro209Leu mutant was also the one affecting the clearance of SOD1_G93A the most. Note that the most severe phenotypes are also associated with the Pro209Leu mutation (Selcen et al. 2009). However, it does not fully explain why this variant is associated with cardial symptoms while the two other variants are more frequently linked to distal myopathy or peripheral neuropathy. In fact, there is even one patient reported with a Pro209Leu variant who only suffers from myofibrillar myopathy but not cardiomyopathy (Andersen et al. 2017). Similarly, the mouse does not seem to recapitulate this cardial phenotype either; as a transgenic knock-in model of a Pro215Leu mutation in BAG3, equivalent to the human Pro209Leu mutant, did not show any abnormal cardial function or morphology up to 16 months of age (Fang et al. 2019). Similar to SQSTM1/p62 mutations, to which BAG3-Pro209 mutants bind stronger, genotype-phenotype correlations thus only poorly predict the clinical presentation (Long et al. 2017). However, the possibility that other modifying or (epi-) genetic factors contribute to clinical differences in both BAG3 and SQSTM1/p62 linked diseases cannot be excluded.

To conclude, despite the distinct phenotypes associated with Pro209 mutations in BAG3, they all seem to induce aggresome formation causing the sequestration of PQC factors. This suggests that if a therapy for one of the Pro209-associated diseases can be identified, it may also be beneficial to other Pro209-associated phenotypes.

## Materials and Methods

### In vitro mutagenesis

Mutations were introduced through site-directed mutagenesis using the wild type BAG3-GSGS-GFP construct in the pEGFP-N1 vector (a kind gift of Josée N. Lavoie). Point mutations were introduced with following primers:

(Pro209Ser):

Fw: CGCGGGGGTACATCTCCATTTCGGTGATACACGAGCAGAA

Rv: TTCTGCTCGTGTATCACCGAAATGGAGATGTACCCCCGCG

(Pro209Leu):

Fw: CGCGGGGGTACATCTCCATTCTGGTGATACACGAGCAGAA

Rv: TTCTGCTCGTGTATCACCAGAATGGAGATGTACCCCCGCG

(Pro209Gln):

Fw: CGCGGGGGTACATCTCCATTCAGGTGATACACGAGCAGAA

Rv: TTCTGCTCGTGTATCACCTGAATGGAGATGTACCCCCGCG

(Glu455Lys):

Fw: AAAAAGTACCTGATGATCAAAGAGTATTTGACCAAAGAGC

Rv: GCTCTTTGGTCAAATACTCTTTGATCATCAGGTACTTTTT

Incorporation of the respective mutations was verified by Sanger sequencing.

### Generation of stable cell lines

HEK293T cells were transduced with lentivirus containing the HSPB8 ORF (NM_014365) in pLENTI6/V5 (Life Technologies, Carlsbad, CA, USA). In brief, HEK293T cells (purchased from ATCC, Teddingtin, Middlesex, UK) were transiently transfected with packaging (pCMV dR8.91), envelope (pMD2-VSV) and pLenti6/V5 plasmids using linear polyethylenimine (PEI) (23966-1, PolySciences Europe, Hirschberg an der Bergstrasse, Germany) or PEI MAX (24765-1, PolySciences Europe, Hirschberg an der Bergstrasse, Germany). After 48 h, the virus containing supernatant was collected, filtered and transferred to fresh HEK293T cells for infection. Positive cells were selected by blasticidine selection. Cells were cultured at 37 °C and 5% CO2 in DMEM (Life Technologies, Carlsbad, CA, USA) supplemented with 10% Fetal Bovine Serum, 1% Glutamine and 1% Penicillin-Streptomycin (Life Technologies, Carlsbad, CA, USA).

### Western blot and Filter retardation assay

For BAG3 solubility assessment in western blot (WB) and filter retardation assays (FRA) (**Fig. 1**), stable transfected HEK293T with HSPB8-V5 were plated at 90,000 cells/well, C2C12 at 65,000 cells/well, NSC-34 at 90,000 cell/well in 12-well. After 24 h stable transfected HEK293T with HSPB8-V5 were transiently transfected using Lipofectamine3000/P3000 reagent with pEGFP-N1 as mock or BAG3-GFP constructs (wild type or mutants) alone or co-transfected with plasmid encoding SOD1_G93A or poly-GA (kindly provided by Prof. Daisuke Ito (Keio University School of Medicine, Tokyo, Japan)), following the manufacturers’ instructions (L3000-001; Life Technologies, Carlsbad, CA, USA). C2C12 cells were transiently transfected using Lipofectamine3000/P3000 reagent, NSC-34 were transiently transfected using Lipofectamine (18324010; Life Technologies, Carlsbad, CA, USA) transferrin (Sigma-Aldrich) with pEGFP-N1 as mock or BAG3-GFP constructs (wild type or mutants). Samples were harvested and centrifuged 5 min at 100 *g* at 4 °C. Cells were resuspended in NP-40 lysis buffer (150 mM NaCl, 20 mM TrisBase, NP-40 0.05%, 1.5 mM MgCl_2_, Glycerol 3%, pH 7.4) added DTT and Complete Protease inhibitor (Roche Applied Science, Indianapolis, IN, USA), and passed through a syringe 10 times. Lysed cells were centrifuged at 16,100 *g* for 15 min. Supernatants were collected and pellets resuspended in the same volume of NP-40 buffer without protease inhibitors and DTT, and finally sonicated. For the evaluation of the effects of BAG3 mutations on its chaperone-activity towards aggregation-prone proteins (SOD1_G93A) (**Fig. 6**), HEK293T cells were co-transfected with BAG3-GFP constructs and SOD1_G93A encoding plasmid, as described above. Cells were then harvested and centrifuged for 5 min at 100 *g* at 4 °C. The pelleted cells were resuspended in PBS with protease inhibitors cocktail (Sigma-Aldrich, Saint Louis, MI, USA) and lysed using slight sonication. SDS-PAGE was performed loading 10 μg of total protein extracts heated to 100 °C for 5 min in sample buffer (0.6 g/100 mL Tris, 2 g/100 mL SDS, 10% glycerol, 5% β-mercaptoethanol, pH 6.8). Proteins were electro-transferred to nitrocellulose membrane (cat. 1620115; Bio-Rad Laboratories, Hercules, CA, USA) using Trans-turbo transfer System (cat. 1704150; Bio-Rad Laboratories, Hercules, CA, USA). Filter Retardation Assay was performed using a Bio-Dot SF Apparatus (Bio-Rad Laboratories, Hercules, CA, USA), as previously described (Sau et al. 2007). A 0.2-μm cellulose acetate membrane (Whatman 100404180) was treated in 20% methanol and washed in PBS. Then 3 μg (for BAG3-GFP) and 6 μg (for SOD1_G93A) of total protein were loaded and filtered by gentle vacuum. An equal amount of protein was loaded for each sample after correcting with a BCA assay. Then FRA membranes were washed in PBS and rinsed in 20% methanol. WB and FRA membranes were treated with a blocking solution of non-fat dried milk (5%) in TBS-Tween (20 mM Tris-HCl pH 7.5, 0.5 M NaCl, 0.05% Tween-20) for 1 h. WB and FRA membranes were then probed using the following antibodies: mouse monoclonal anti-GFP antibody (ab1218, Abcam, Cambridge, UK), mouse monoclonal anti-α-tubulin (T6199, Sigma-Aldrich, Saint Louis, MI, USA), homemade rabbit polyclonal anti-BAG3 (Carra et al. 2008), homemade rabbit polyclonal anti-HSPB8 (#23 (Carra et al. 2005)), mouse monoclonal anti-HSPB8 (ab66063, Abcam, Cambridge, UK), rabbit polyclonal anti-Cu/Zn superoxide dismutase SOD1 (SOD-100, Enzo Life Sciences, Farmingdale, NY, USA), anti-ubiquitin (3936, Cell Signaling Technology, Danvers, MA, USA), rabbit polyclonal anti-BAG3 (10599-1-AP, Proteintech, Rosemont, IL, USA). Membranes were then washed 3 times for 10 min. Immunoreactivity was detected using the following secondary peroxidase conjugated antibodies: goat anti-rabbit and anti-mouse IgG-HRP (111-035-003, 115-035-003; Jackson ImmunoResearch Laboratories, Inc. Cambridge, UK) and enhanced chemiluminescent (ECL) detection reagent (Clarity™ ECL western blotting substrate, cat. 1705060; Bio-Rad Laboratories, Hercules, CA, USA). Images were acquired using a Chemidoc XRS System (Bio-Rad Laboratories, Hercules, CA, USA) and optical intensity was analysed using Image Lab Software (Bio-Rad Laboratories, Hercules, CA, USA).

### Flow Cytometric analysis of inclusions (FloIT)

HEK293T cells that were lentiviral transduced with HSPB8-V5 and were plated in 24-well plates at 75,000 cells/well. After 24 h, cells were transiently transfected using Lipofectamine3000/P3000 reagent, as previously described. After 48 h, medium was removed and cells were harvested in PBS with 10% FBS (Gibco, Thermo Fisher Scientific, Waltham, MA, USA) and centrifuged for 5 min at 100 *g* at 4 °C. Cells were resuspended in PBS with 10% FBS (Gibco, Thermo Fisher Scientific, Waltham, MA, USA) and an aliquot was analyzed by flow cytometry to determine the transfection efficiency in respect to untransfected control cells. Flow cytometry was performed using NovoCyte Flow Cytometer 3000 (ACEA Biosciences Inc., Agilent, Santa Clara, CA, USA) and results were analysed by NovoExpress software 1.2.5 (ACEA Biosciences Inc., Agilent, Santa Clara, CA, USA). For transfection efficiency analysis, excitation wavelengths and emission collection windows were FITC (488 nm, 530/50 nm), Pacific Blue (405 nm, 445/45 nm). Subsequently, a solution of PBS containing 1% (v/v) Triton X-100, a cocktail of Protease inhibitors (Sigma-Aldrich, Saint Louis, MI, USA) and DAPI (0.02 ug/ul) was added to a final concentration of 0.5% (v/v) Triton X-100 and DAPI 0.01 μg/μl. After two minutes incubation at room temperature, the cell lysates were analysed by flow cytometry. Three untransfected control samples without DAPI were analysed to set gates on nuclei population. Voltage of 418 (FSC), 199 (SSC), 373/482 (FITC for cell transfection or inclusion analysis respectively), 501 (Pacific Blue) were used. Nuclei were counted based on the Pacific Blue positive population. Inclusions were identified for fluorescence and FSC compared to cells transfected with eGFPN1 vector as control. Following the equation set by Whiten et al. (2016) the number of inclusions was normalized to the number of counted nuclei and reported as inclusions/100 transfected cells. Nuclei population was analysed based on FITC fluorescence and a percentage of nuclei enriched with GFP-positive particles was determined.

### Protein solubility predictions

To predict protein solubility, we used the CamSol browser (accessed on September 2^nd^ 2019; http://www-vendruscolo.ch.cam.ac.uk/camsolmethod.html) and Tango (accessed on September 3^rd^ 2019; http://tango.crg.es). As input for the CamSol method, we either inserted the full-length protein sequence of BAG3 (NP_004272.2) (**Fig. S2**) or the 20 amino acids surrounding the second IPV-motif (HQLPRGYlSI**<u>P</u>**VIHEQNVTRP; **Fig. 1d**). In case of the latter, we also performed the solubility calculations for each of the respective mutants by replacing the Pro209 by either Ser, Leu or Gln. We used the CamSol Intrinsic method, as described in Sormanni et al. 2015.

For Tango, we inserted a protein sequence of 70 amino acids spanning the second IPV-motif (SQSPAASDCSSSSSSASLPSSGRSSLGSHQLPRGYISI**<u>P</u>**VIHEQNVTRP AAQPSFHQAQKTHYPAQQGEY)(**Fig. 1e**). The parameters were as following: no protection at the N-terminus or C-terminus of the peptide sequence, pH was selected as 7, temperature 298.15 K, ionic strength of 0.02M, and a concentration of 1M. We selected and plotted Beta-aggregation for both the wild type sequence as the three IPV-mutants (Ser/Leu/Gln).

### Co-immunoprecipitation

HEK293T stable cell lines for HSPB8-V5 were transiently transfected with different wild type or mutant BAG3-GFP constructs using PEI (23966-1, PolySciences Europe, Hirschberg an der Bergstrasse, Germany). Or alternatively, the reverse experiment was performed by transiently transfecting HeLa cells. After 48 h, cells were lysed with lysis buffer [20mM Tris-HCl pH 7.4, 2.5 mM MgCl_2_, 100 mM KCl, 0.5% Nonidet P-40, Complete Protease inhibitor (Roche Applied Science, Indianapolis, IN, USA)] and incubated on ice for 30 min. Samples were centrifuged for 10 min at 20,000 *g* and equal amounts of supernatant (NP40-soluble fraction only) was loaded on GFP-Trap beads (gta-20, Chromotek, Martinsried, Germany). Beads were incubated with the protein lysate for 1 h at 4 °C and washed three times with wash buffer [20 mM Tris-HCl pH 7.4, 2.5 mM MgCl_2_, 100 mM KCl, Complete Protease inhibitor (Roche Applied Science, Indianapolis, IN, USA)]. Proteins were eluted from the beads with Sarkosyl elution buffer (140 mM NaCl, 50 mM Tris-HCl pH 8, 1 mM EDTA, 0.3% Sarkosyl, 10% glycerol) before being supplemented with NuPAGE LDS sample buffer (Life Technologies, Carlsbad, CA, USA) and loaded on 4-12% NuPAGE gels (Life Technologies, Carlsbad, CA, USA). Proteins were transferred to nitrocellulose membranes (Hybond-P; GE Healthcare, Wauwatosa, WI, USA) and decorated with antibodies against GFP (ab290, Abcam, Cambridge, UK), V5 (R96025, Invitrogen, Carlsbad, CA, USA), SQSTM1/p62 (5114, Cell Signaling, Danvers, MA, USA), Hsp70/Hsc70 (ab5439, Abcam, Cambridge, UK), or Tubulin (ab7291, Abcam, Cambridge, UK). Samples were detected using enhanced chemiluminescent ECL Plus (Pierce, Life Technologies, Carlsbad, CA, USA) and LAS4000 (GE Healthcare, Wauwatosa, WI, USA).

For the reverse experiment, co-immunoprecipitation of BAG3 after HSPB8 pull-down (**Fig. S7**) was performed as previously described in Minoia et al. 2014. In brief, HeLa cells were transfected using Lipofectamine 2000 reagent (Invitrogen, Carlsbad, CA, USA) with empty vector or BAG3-GFP constructs (wild type or mutants), according to manufacturer’s instructions. 24 h post-transfection cells were lysed in lysis buffer (150 mM NaCl, 0.5% NP40, 1.5 mM MgCl_2_, 20 mM Tris-HCl pH 7.4, 3% glycerol, 1 mM DTT, Complete Protease inhibitor (Roche Applied Science, Indianapolis, IN, USA)). The cell lysates were centrifuged and cleared with A/G beads (Santa Cruz Biotechnology, Inc., Santa Cruz, CA, USA) at 4 °C for 1 h. Rabbit TrueBlot beads (Tebu-bio) were incubated at 4 °C for 1 h with home-made rabbit HSPB8 antibody (Carra et al. 2005) or with rabbit serum (NRS), used as a control. Rabbit TrueBlot beads complexed with the specific antibodies were added to the precleared lysates. After incubation for 1 h at 4 °C, the immune complexes were centrifuged. Beads were washed four times with the lysis buffer; both co-immunoprecipitated proteins and input fractions were resolved on SDS-PAGE followed by western blot.

### Fluorescence microscopy and immunofluorescence

For quantification of protein aggregates, HEK293T stable cell line for HSPB8-V5, NSC-34 or C2C12 cells were plated in 24-well plates containing poly-D lysine (P-7280, Sigma-Aldrich, Saint Louis, MI, USA) coated coverslips and then transfected with wild type or mutant BAG3-GFP constructs as described above for WB and FRA experiments. For protein aggregation-prone behaviour evaluation, C2C12 were plated at 50,000 cells/ml, NSC-34 at 70,000 cell/ml in 24-well. Cells were then fixed using a 1:1 solution of 4% paraformaldehyde (PFA) and 4% sucrose in 0.2 N PB (0.06 M KH_2_PO_4_, 0.31 M Na_2_HPO_4_; pH 7.4) for 25 min at 37 °C. Nuclei were stained with Hoechst (1:2000 in PBS; 33342, Sigma-Aldrich, Saint Louis, MI, USA). Images were captured by Axiovert 200 microscope (Zeiss, Oberkochen, Germany) with a photometric CoolSnap CCD camera (Ropper Scientific, Trenton, NJ, USA). Images were processed using Metamorph software (Universal Imaging, Downingtown, PA). Six different fields were captured for each sample, of which each field of view contained an average of 95 cells. This summed up to a total of WT=687 cells, Pro209Ser=546 cells, Pro209Leu=473 cells, Pro209Gln=651 cells, Glu455Lys=499 cells.

For colocalization studies of BAG3 and vimentin, the same HEK293T stable cell line for HSPB8-V5 were grown on poly-D-lysine (P7280-5×5MG, Sigma-Aldrich, Saint Louis, MI, USA) coated glass coverslips. Cells were transfected using PEI MAX (24765-1, PolySciences Europe, Hirschberg an der Bergstrasse, Germany) for wild type or mutant BAG3-GFP and fixed in ice-cold methanol (67-65-1, Sigma-Aldrich, Saint Louis, MI, USA) for 20 minutes. After blocking with 5% BSA (9048-46-8, Sigma-Aldrich, Saint Louis, MI, USA), cells were incubated with anti-GFP (Alexa Fluor 488 anti-GFP antibody; 338008, BioLegend, San Diego, CA, USA) and anti-vimentin (dilution 1:200; ab28028, Abcam, Cambridge, UK) for one hour at room temperature. After secondary antibody incubation, nuclei were stained with Hoechst33342 (H3570, Life Technologies, Carlsbad, CA, USA) and cells were mounted with DAKO fluorescent mounting medium (S3023, DAKO). Imaging was performed on a Zeiss LSM700 laser scanning confocal microscope using a 63x/1.4 NA objective. Image analysis was done in ImageJ/FIJI (Schindelin et al. 2012, Schneider et al. 2012).

For colocalization between BAG3 and SQSTM1/p62 or FLAG-HDAC6, HEK293T stable cell line for HSPB8-V5 were grown on polylysine-coated glass coverslip. Cells were then transfected with cDNAs encoding for wild type or mutant BAG3-GFP alone or, when indicated, FLAG-HDAC6 (Fischle W. et al. 1999). After washing with cold PBS, cells were fixed with 3.7% formaldehyde in PBS for 9 minutes at room temperature, followed by permeabilization with cold acetone for 5 minutes at −20 °C. After blocking for 1 h at room temperature with BSA 3% and 0.1% Triton X-100, cells were incubated with anti-SQSTM1/p62 (sc-28359, Santa Cruz Biotechnology, Inc., Santa Cruz, CA, USA) or anti-FLAG (F3165, Sigma-Aldrich, Saint Louis, MI, USA) overnight at 4 °C. Cells were washed and incubated for 1 h at room temperature with a mouse secondary antibody (A21203, Life Technologies, Carlsbad, CA, USA); nuclei were stained with DAPI. Images were obtained using a Leica SP8 AOBS system (Leica Microsystems) and a 63x oil-immersion lens. Image analysis was done in ImageJ/FIJI (Schindelin et al. 2012, Schneider et al. 2012).

For aggresome assessment after dynein inhibition, HEK293T stable cell line for HSPB8-V5 were grown on poly-D-lysine (P7280-5×5MG, Sigma-Aldrich, Saint Louis, MI, USA) coated glass coverslips. Cells were transfected using PEI MAX (24765-1, PolySciences Europe, Hirschberg an der Bergstrasse, Germany) for wild type or mutant BAG3-GFP and subjected to 6 h dynein inhibition with Dynarrestin (25μM, SML2332-5MG, Sigma-Aldrich, Saint Louis, MI, USA) or erythro-9-(2-hydroxy-3-nonyl)adenine (EHNA; 100μM, E114-25MG, Sigma-Aldrich, Saint Louis, MI, USA) prior to fixation with 2% PFA (28906, Life Technologies, Carlsbad, CA, USA) for 20 minutes at room temperature. After permeabilization for two minutes with 0.1% Triton X-100 (9002-93-1, Sigma-Aldrich, Saint Louis, MI, USA), cells were first blocked with 5% BSA (9048-46-8, Sigma-Aldrich, Saint Louis, MI, USA) and then incubated with GFP-booster Atto488 (gba488-100, Chromotek, Martinsried, Germany) for one hour at room temperature. Nuclei were stained with Hoechst33342 (H3570, Life Technologies, Carlsbad, CA, USA) and cells were mounted with DAKO fluorescent mounting medium (S3023, DAKO). Imaging was performed on a Zeiss LSM700 laser scanning confocal microscope using a 63x/1.4 NA objective. Image analysis was done in ImageJ/FIJI (Schindelin et al. 2012, Schneider et al. 2012).

### Fluorescence recovery after photobleaching (FRAP) assay

HeLa cells were transfected with BAG3-GFP wild type or mutant constructs and with either P62-mCherry and mScarlet-HSP70 constructs and imaged 48 hours after transfection in a μ-slide 8-well (80826, Ibidi, Martinsried, Germany) in FluoroBrite DMEM medium (Life Technologies, Carlsbad, CA, USA) supplemented with 10% fetal bovine serum and 4mM L-glutamine at 37°C and 5% CO2. FRAP measurements were performed on a Zeiss LSM700 laser scanning confocal microscope using a PlanApochromat 63x/1.4 NA objective.

Image sequences (512 x 512 pixels, 117 nm/pixel) were acquired at 1 frame per 3 sec (Bag3-GFP and mScarlet-HSP70 FRAP) or 1 frame per 2 sec (P62-mCherry FRAP) for the duration of the experiments, as indicated in the figures. Three to five pre-bleach sequences preceded photobleaching in a 70 x 70 pixel region at 100% of a 5mW 488nm laser (Bag3-GFP FRAP) or 100% of a 10 mW 555nm (P62-mCherry and mScarlet-HSP70 FRAP) for 2 sec. FRAP sequences were recorded from 6 cells per genotype and intensities in the bleached region were measured with ImageJ (Schindelin et al. 2012, Schneider et al. 2012) and plotted over time. Imaging and photobleaching settings were kept identical for all wild type and mutant Bag3 cells within the three different FRAP experiments.

### Live-cell time-lapse imaging

HEK293T cells or HeLa cells were transfected with GFP-tagged BAG3 wild type or mutant constructs and imaged once per hour using an IncuCyte S3 instrument (Essen BioScience, UK). In case of co-localization experiments, cells were co-transfected with mCherry-tagged SQSTM1/p62 constructs (a kind gift from Prof. Sascha Martens (Max F. Perutz Laboratories, University of Vienna, Vienna Biocenter, Vienna, Austria)) or mScarlet-tagged Hsp70 constructs (we cloned Hsp70 into the pmScarlet_C1 plasmid, which was a kind gift from Dorus Gadella and available from Addgene (#85042)). Images were taken with a 20x objective at 37°C and 5% CO_2_. GFP-tagged proteins were excited by a green laser for 300 ms and with a red laser for red-fluorescently tagged proteins for 400 ms. Images were exported and further analyzed in ImageJ (Schindelin et al. 2012, Schneider et al. 2012).

### Cycloheximide treatment

To determine protein turnover, a cycloheximide wash-out experiment was performed on HEK293T cells that stably overexpress HSPB8-V5. Cells were transiently transfected using PEI MAX (24765-1, PolySciences Europe, Hirschberg an der Bergstrasse, Germany) with wild type or mutant BAG3-GFP constructs. One day after transfection, cells were subjected to cycloheximide treatment (100 μg/ml) for either 6 h or 12 h. As a control, cells were subjected to dimethyl sulfoxide (DMSO) for 12 h. After the treatment, cells were collected in ice-cold PBS and proteins were extracted with RIPA buffer [1% Nonidet P-40, 150 mM NaCl, 0.1% SDS, 0.5% deoxycholic acid, 1 mM EDTA, 50 mM Tris-HCl pH 7.5, cOmplete Protease Inhibitor Cocktail (Roche Applied Science, Indianapolis, IN, USA), Phospho-STOP inhibitor mix (05 892 970 001, Roche Applied Science, Indianapolis, IN, USA)]. The soluble and insoluble fraction were separated by centrifugation at max speed for 10 minutes at 4 °C. Insoluble pellets were resuspended in, while the soluble supernatant was subjected to a BCA assay (23225, Pierce BCA Protein Assay Kit) for protein quantification. Equal amounts of protein were loaded on 4-12% BisTris NuPAGE gels (Life Technologies, Carlsbad, CA, USA). Proteins were transferred to nitrocellulose membranes (Hybond-P; GE Healthcare, Wauwatosa, WI, USA). Ponceau S was used to assess equal loading of the insoluble fraction. Membranes were further decorated with antibodies against GFP (ab290, Abcam, Cambridge, UK), and β-Actin/ACTB (A5316, Sigma-Aldrich, Saint Louis, MI, USA). Samples were detected using enhanced chemiluminescent ECL Plus (Pierce, Life Technologies, Carlsbad, CA, USA) and LAS4000 (GE Healthcare, Wauwatosa, WI, USA). Protein densitometry was performed using ImageJ/FIJI (Schindelin et al. 2012, Schneider et al. 2012) and plotted over time relative to time-point zero.

### Autophagy assay

For western blot analysis of autophagic flux, HEK293T cells stably transduced with HSPB8 were transiently transfected using PEI MAX (24765-1, PolySciences Europe, Hirschberg an der Bergstrasse, Germany) for wild type or mutant BAG3-GFP. After 24 h, cells were cultured in serum-deprived medium with bafilomycin A1 (10 nM) for two hours. Proteins were extracted with RIPA buffer [1% Nonidet P-40, 150 mM NaCl, 0.1% SDS, 0.5% deoxycholic acid, 1 mM EDTA, 50 mM Tris-HCl pH 7.5, cOmplete Protease Inhibitor Cocktail (Roche Applied Science, Indianapolis, IN, USA), Phospho-STOP inhibitor mix (05 892 970 001, Roche Applied Science, Indianapolis, IN, USA)], quantified using BCA (23225, Pierce BCA Protein Assay Kit). Equal amounts of protein were then loaded on 12% NuPAGE gels (Life Technologies, Carlsbad, CA, USA). Proteins were transferred to nitrocellulose membranes (Hybond-P; GE Healthcare, Wauwatosa, WI, USA) and decorated with antibodies against GFP (ab290, Abcam, Cambridge, UK), LC3B (L7543, Sigma-Aldrich, Saint Louis, MI, USA), and β-Actin/ACTB (A5316, Sigma-Aldrich, Saint Louis, MI, USA). Samples were detected using enhanced chemiluminescent ECL Plus (Pierce, Life Technologies, Carlsbad, CA, USA) and LAS4000 (GE Healthcare, Wauwatosa, WI, USA).

### Statistics

Statistical analyses have been performed using the statistical tests as stated in each figure legend. This comprised Student T tests or One-Way ANOVA with Bonferroni’s multiple comparisons tests. The statistical analysis was performed using PRISM software (GraphPad Software, La Jolla, CA, USA).

## Acknowledgments

This research was a collaboration between the Universities of Antwerp, Modena and Milan. The project was in part funded by the Fund for Scientific Research (FWO-Flanders, to V.T), the Medical Foundation Queen Elisabeth (GSKE, to V.T.), the “Association Belge contre les Maladies Neuromusculaires” (ABMM, to V.T. and E.A.), the Rotary ‘Hope in Head’ program (to E.A. and V.T.), the Association Française contre les Myopathies (AFM, to V.T.), the Muscular Dystrophy Association (MDA, to V.T.) and the European Union’s H2020 grant Solve-RD, ‘Solving the unsolved rare diseases’ under grant agreement 2017-779257 (to VT). This research was also supported by the Fondazione Cariplo, Italy (2014-0686 to A.P. and S.C. and 2017-0747 to V.C.), the Fondazione AriSLA, Italy (ALS_Granulopathy to S.C. and A.P.), the Università degli Studi di Milano e piano di sviluppo UNIMI - linea B (to V.C.), the Fondazione Regionale per la Ricerca Biomedica (FRRB) (Regione Lombardia, TRANS_ALS, project. nr. 2015-0023), Italy (to A.P.), the Research funding program from the University of Modena and Reggio Emilia (FAR2016 to S.C.), the Italian Ministry of University and Research (MIUR), PRIN - Progetti di ricerca di interesse nazionale (2015LFPNMN to A.P. and S.C.), the EU Joint Programme - Neurodegenerative Disease Research (JPND) project. The project is supported through the following funding organisations under the aegis of JPND - www.jpnd.eu. Finally, support was received from the European Union’s Horizon 2020 research and innovation programme under grant agreement N° 643417 (Grant ID: 01ED1601A, CureALS to A.P. and S.C.). We thank Dr. Jonathan Vinet and Centro Interdipartimentale Grandi Strumenti/CIGS (www.cigs.unimo.it) for technical support with microscopy studies and we thank the Finkelstein lab (University of Texas at Austin) for sharing their BioRxiv template.

## Author contributions

EA, BT, LM, BA, VC, FA, SC, AP, and VT designed the experiments. EA, BT, LM, BA, VC, FA, SC, AP, and VT performed the research. EA, BT, LM, BA, VC, FA, SC, AP, and VT analyzed the data. EA, SC, AP, and VT wrote the paper with assistance from all authors. All authors reviewed the manuscript.

## Supplementary Figures

**Fig. S1.**
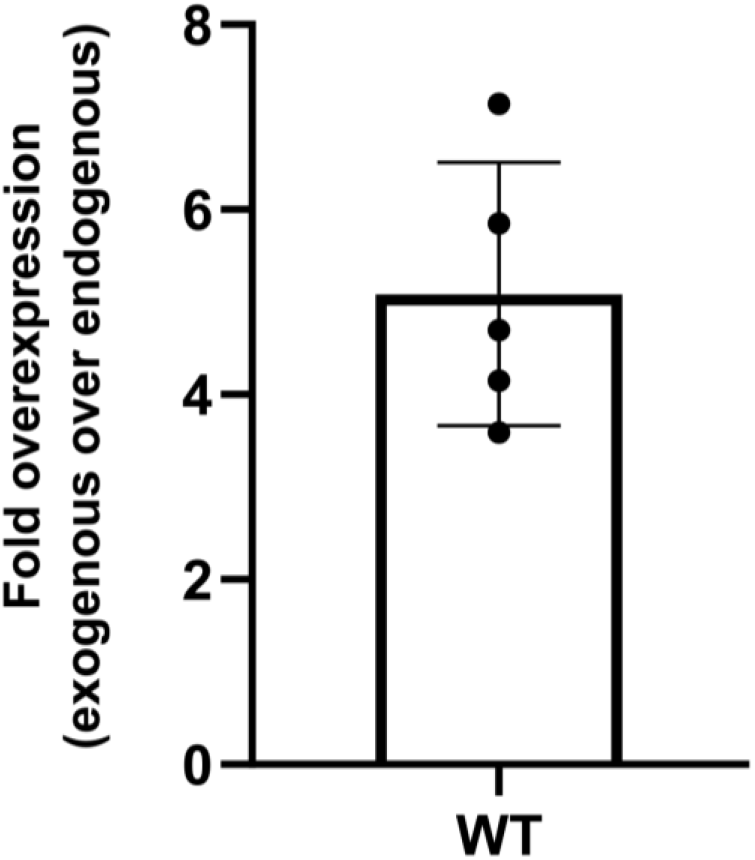
Quantification of the relative fold overexpression of the exogenously GFP-tagged BAG3 over endogenously untagged BAG3. HEK293T-HSPB8-V5 cells were transiently transfected with GFP-tagged wild type BAG3 and protein lysates were analysed by western blot. With an anti-BAG3 antibody both the endogenous and exogenous BAG3 was visualized and the relative fold overexpression was determined with densitometric analysis of the western blot bands. (n=5)

**Fig. S2.**
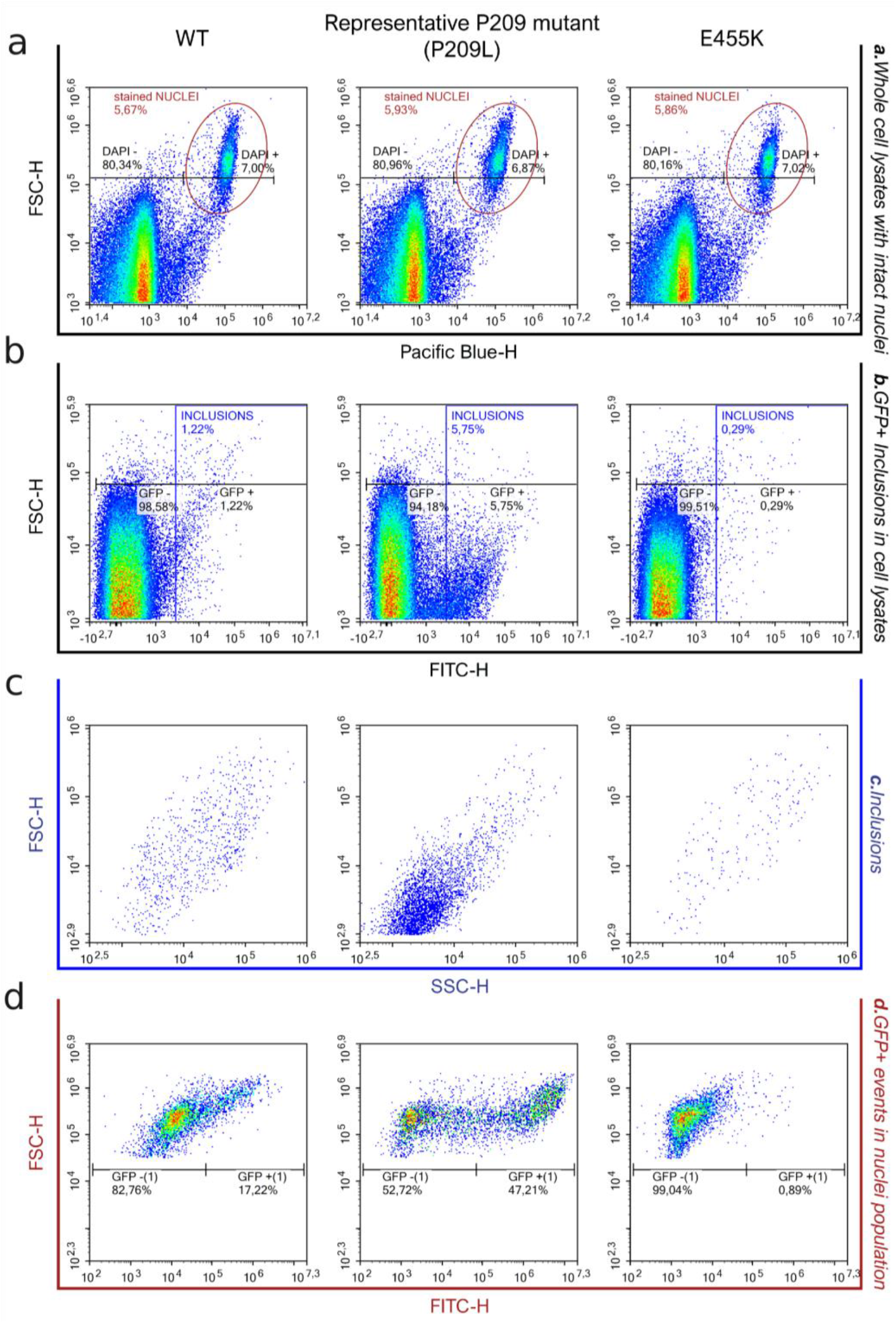
Flow cytometric analysis of inclusions (FloIT) analysis for BAG3-GFP inclusions detection. HEK293T-HSPB8-V5 cells untransfected or transiently transfected with BAG3-GFP wild type or mutants were analysed by flow cytometry for transfection efficiency evaluation (γ = GFP+/cells). (**a**) After plasma membrane lysis, cellular lysates were read: nuclei populations were defined as Pacific Blue + populations after adding DAPI and excluded from cytoplasmic inclusion analysis. (**b**) The remaining particles were defined as GFP+ inclusions in respect to cells transfected with pEGFP-N1. (**c**) Distribution of GFP+ inclusions based on dimension and complexity. (**d**) Stained nuclei were analysed based on FITC fluorescence and dimension.

**Fig. S3.**
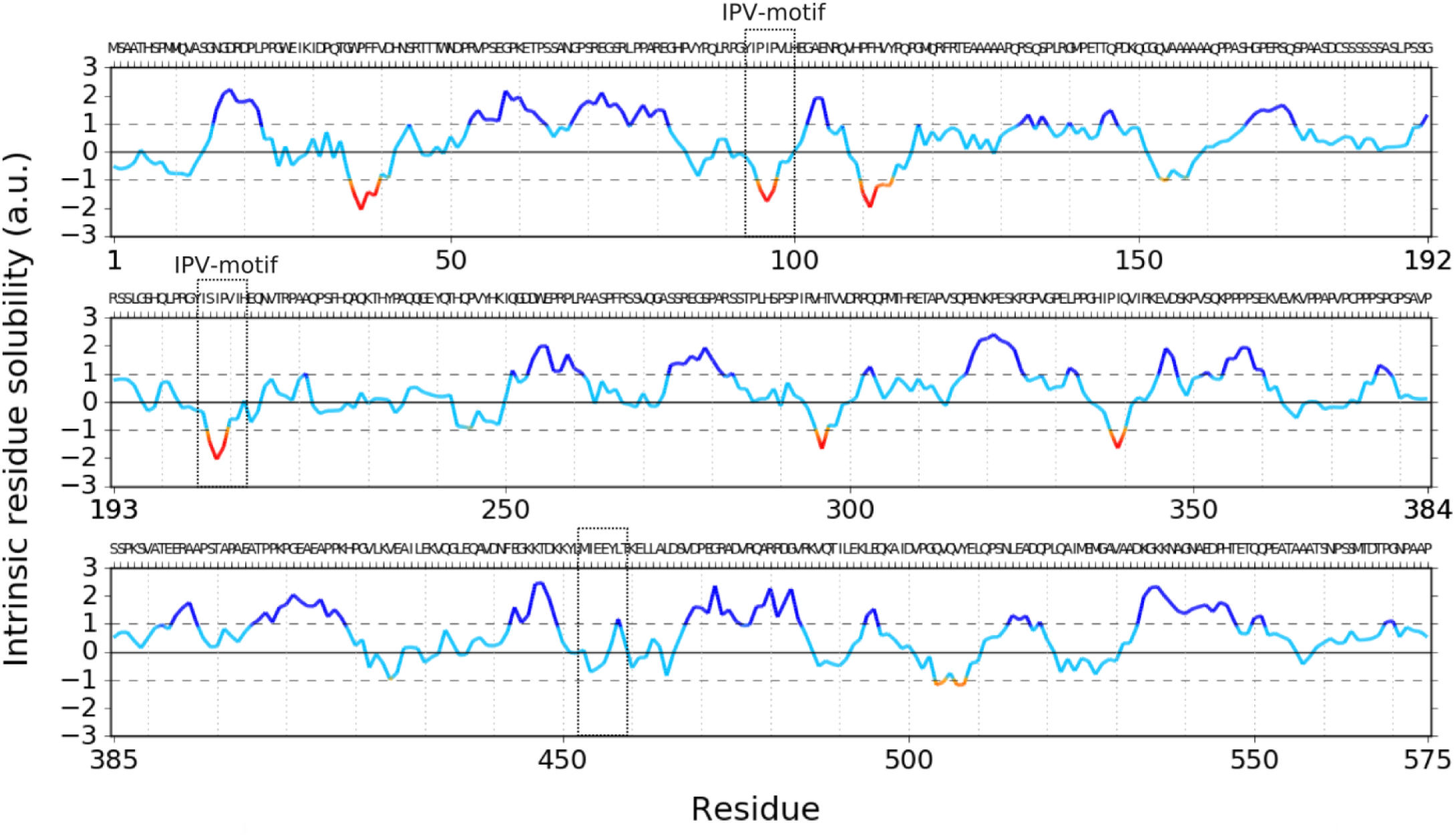
Predicted residue solubility of full length BAG3. Bio-informatic solubility analysis of wild type BAG3 with CamSol software. As input we provided the first 575 amino acids of BAG3. The presented panels are the output as provided by the CamSol software and represent the predicted residue solubility. We highlighted the two IPV-motifs of BAG3 and the location of residue E455 in the BAG-domain.

**Fig. S4.**
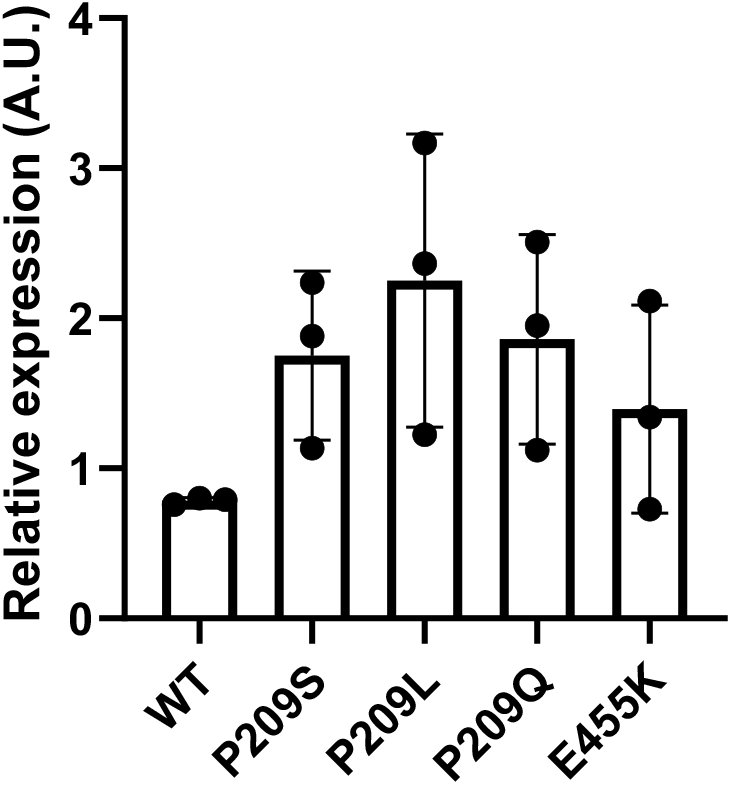
Quantification of total amount of BAG3 wild type or mutants. HEK293T cells that stably overexpress HSPB8-V5 were transiently transfected with wild type or mutant BAG3-GFP constructs. Cells were collected in NP-40 containing buffer and the soluble fraction was separated from the NP-40 insoluble fraction. Both fractions were analysed by western blot and quantified using densitometric analysis. Samples were quantified relative to the wild type (WT). (n=3)

**Fig. S5.**
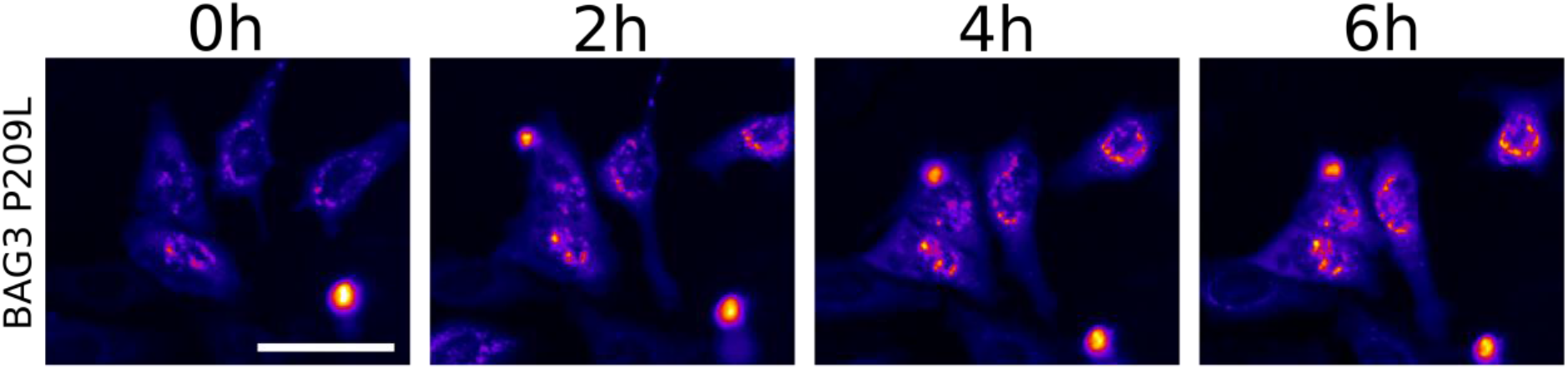
Live-cell time-lapse imaging of GFP-tagged BAG3-Pro209Leu in HeLa cells. HeLa cells were transiently transfected with mutant BAG3-GFP constructs and imaged once per hour. Scale bar = 50 μm.

**Fig. S6.**
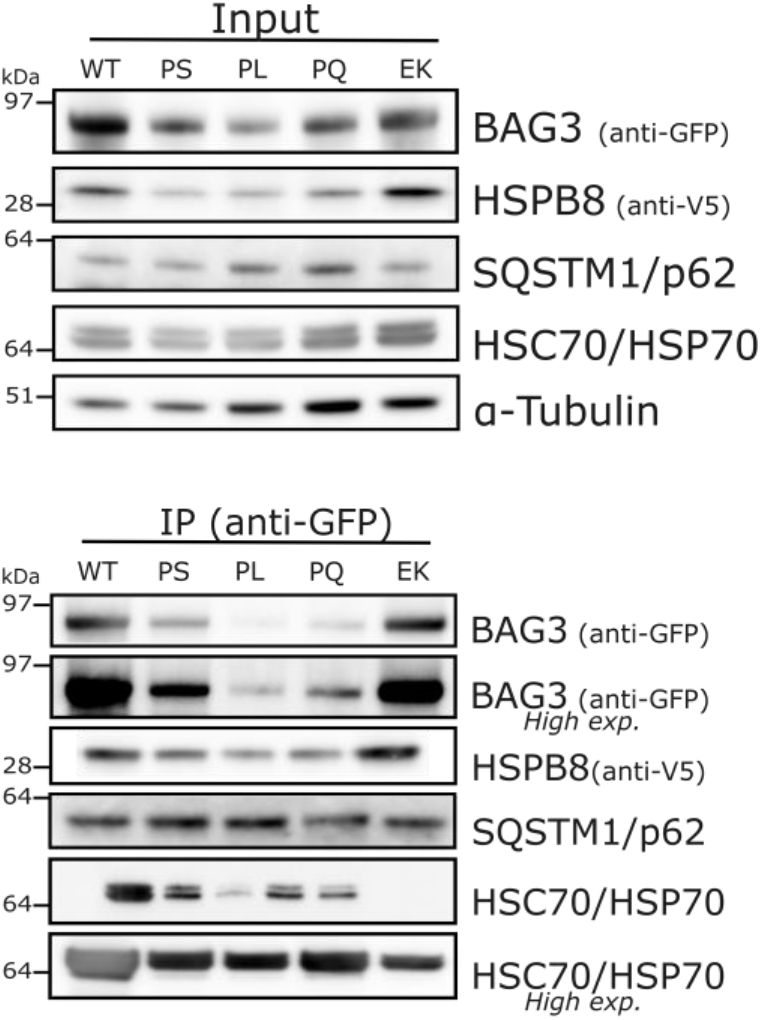
Representative blots from the quantification shown in Figure 4a. HEK293T cells that stably overexpress HSPB8-V5 were transiently transfected with wild type or mutant BAG3-GFP constructs to assess the interaction between BAG3 and components of the CASA-complex. Co-immunoprecipitation of BAG3-GFP and the CASA-complex using the GFP-trap system. Both input and co-immunoprecipitating fraction is displayed. The wild type (WT) or mutants were abbreviated as followed: Pro209Ser (PS), Pro209Leu (PL), Pro209Gln (PQ), Glu455Lys (EK). The amount of interacting proteins was quantified and is displayed in Figure 4a.

**Fig. S7.**
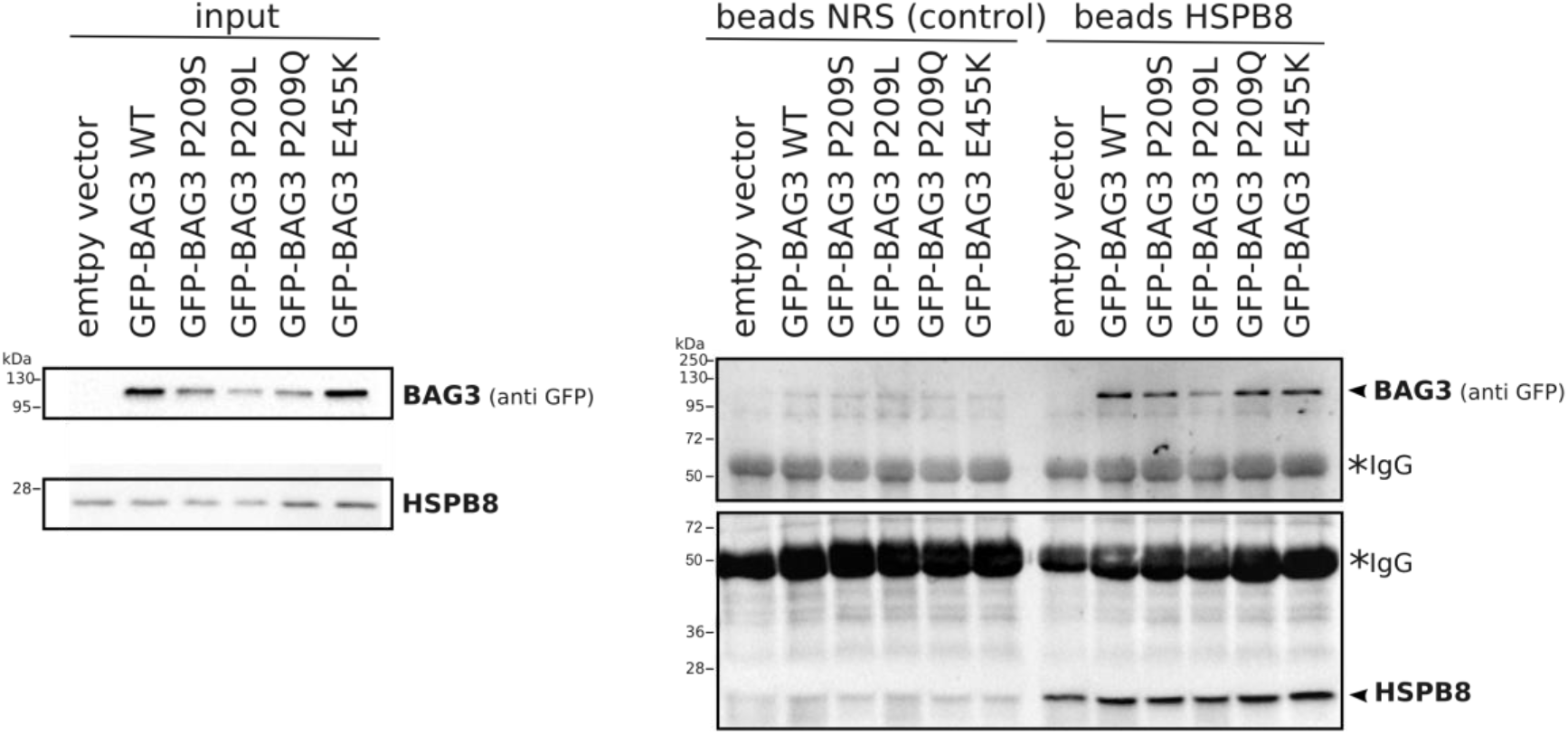
Reverse co-immunoprecipitation of BAG3 after HSPB8 pull-down. HeLa cells were transiently transfected with wild type or mutant BAG3-GFP constructs. Beads coated with a specific anti-HSPB8 antibody or normal rabbit serum (NRS), used as negative control, were mixed with HeLa whole cell lysate. Both input and co-immunoprecipitation samples were loaded on SDS-PAGE. *IgG represents signal from the heavy chains. Arrowheads indicate the correct molecular weight for BAG3-GFP and HSPB8 respectively.

**Fig. S8.**
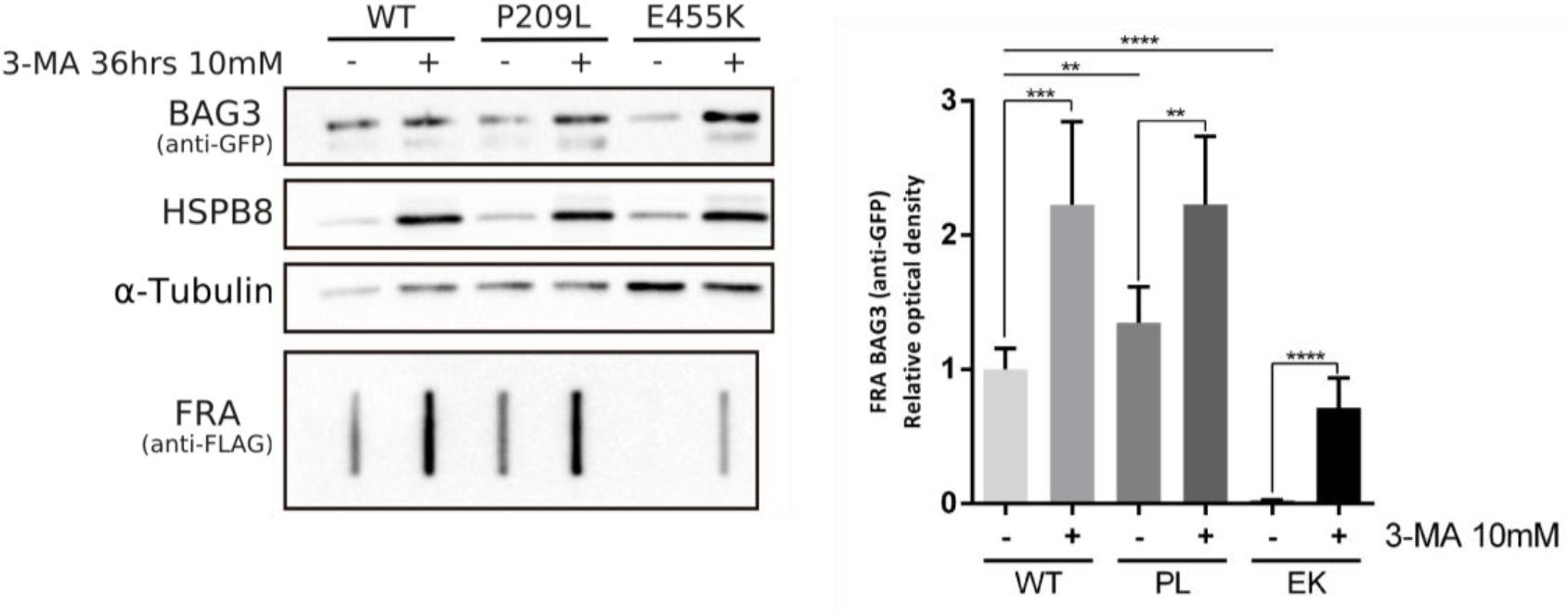
Clearance assay of wild type and mutant BAG3 is prohibited by inhibition of autophagy. HEK293T cells that stably overexpress HSPB8-V5 were transiently transfected with mutant BAG3-GFP constructs. Cells were left untreated or treated with 3-MA for 36 hours after which protein lysates were collected and analysed by western blot and filter retardation assay (FRA). The FRA analysis is displayed for the NP-40 insoluble fraction. Relative optical densities are reported in the graph as means ± SD of normalized values. Student T tests were used for statistical analysis to compare the treated with untreated condition for each respective BAG3 variant (n=3).

**Fig. S9.**
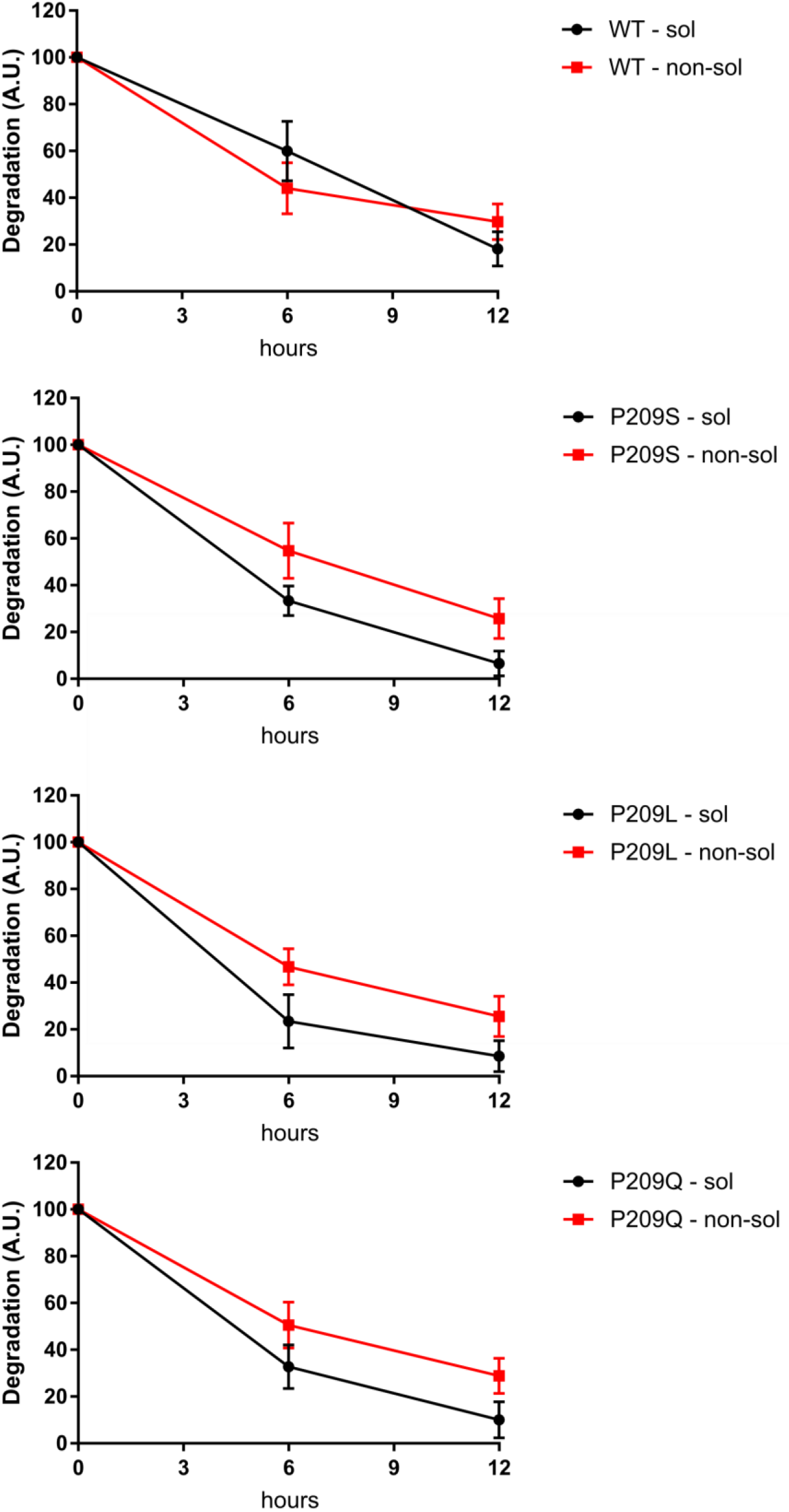
Protein degradation rate soluble versus nonsoluble. The same protein turnover results from Figure 3b are presented here as soluble versus nonsoluble for each of the respective genotypes. The wild type (WT) or mutants were abbreviated as followed: Pro209Ser (PS), Pro209Leu (PL), Pro209Gln (PQ). Data are presented as mean±SD. (n=3)

**Fig. S10.**
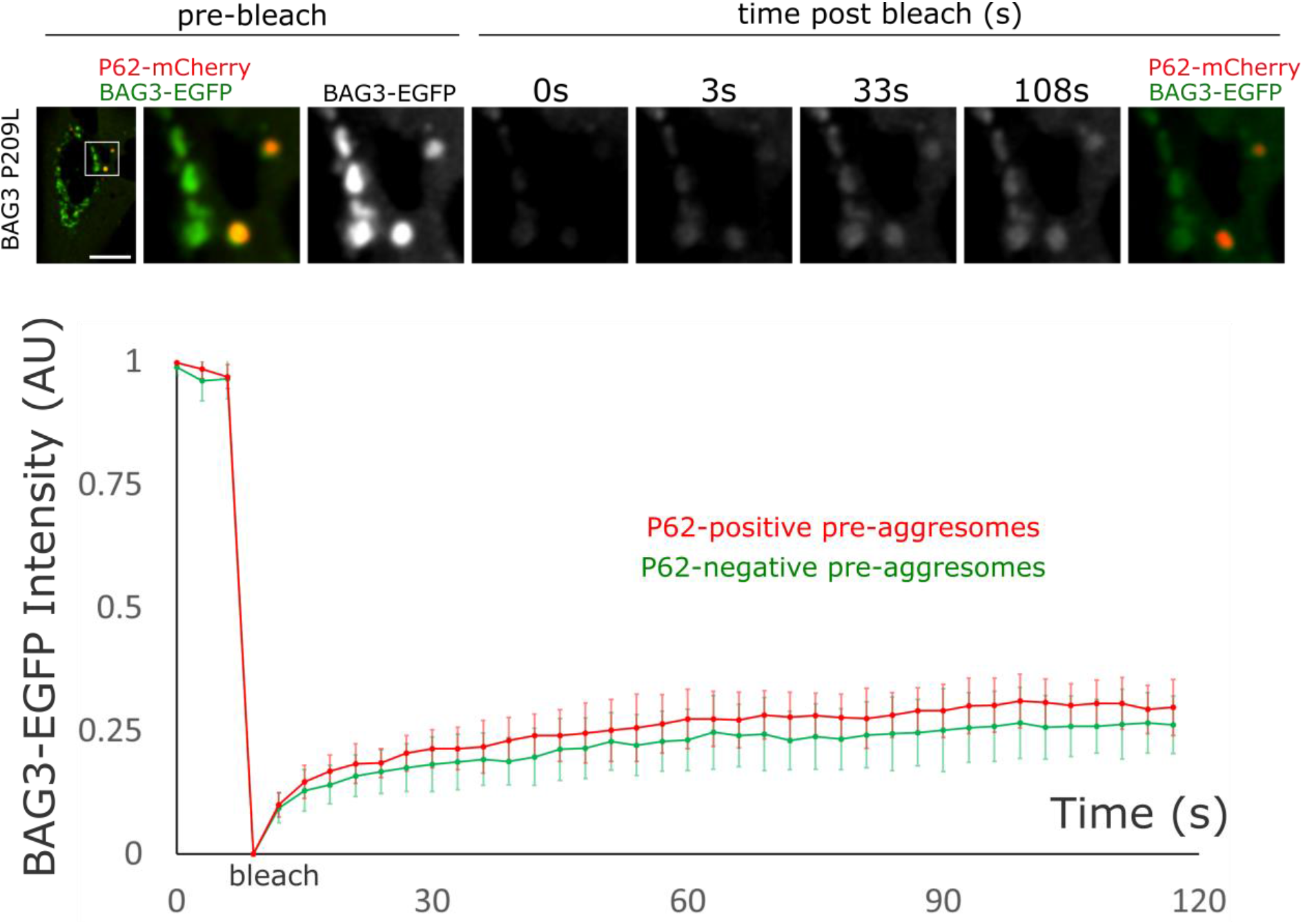
The mobility of BAG3_Pro209 mutants is indifferent in SQSTM1/p62-positive versus SQSTM1/p62-negative pre-aggresome bodies. Fluorescence recovery after photobleaching (FRAP) analysis was performed on HeLa cells that were transiently transfected with BAG3-GFP and SQSTM1/p62-mCherry constructs. Quantification of the fluorescence intensity was plotted over time for mutant BAG3_Pro209Leu cells. Graph bar shows the means (± SD) over time (n=4). Scale bar = 10 μm

**Fig. S11.**
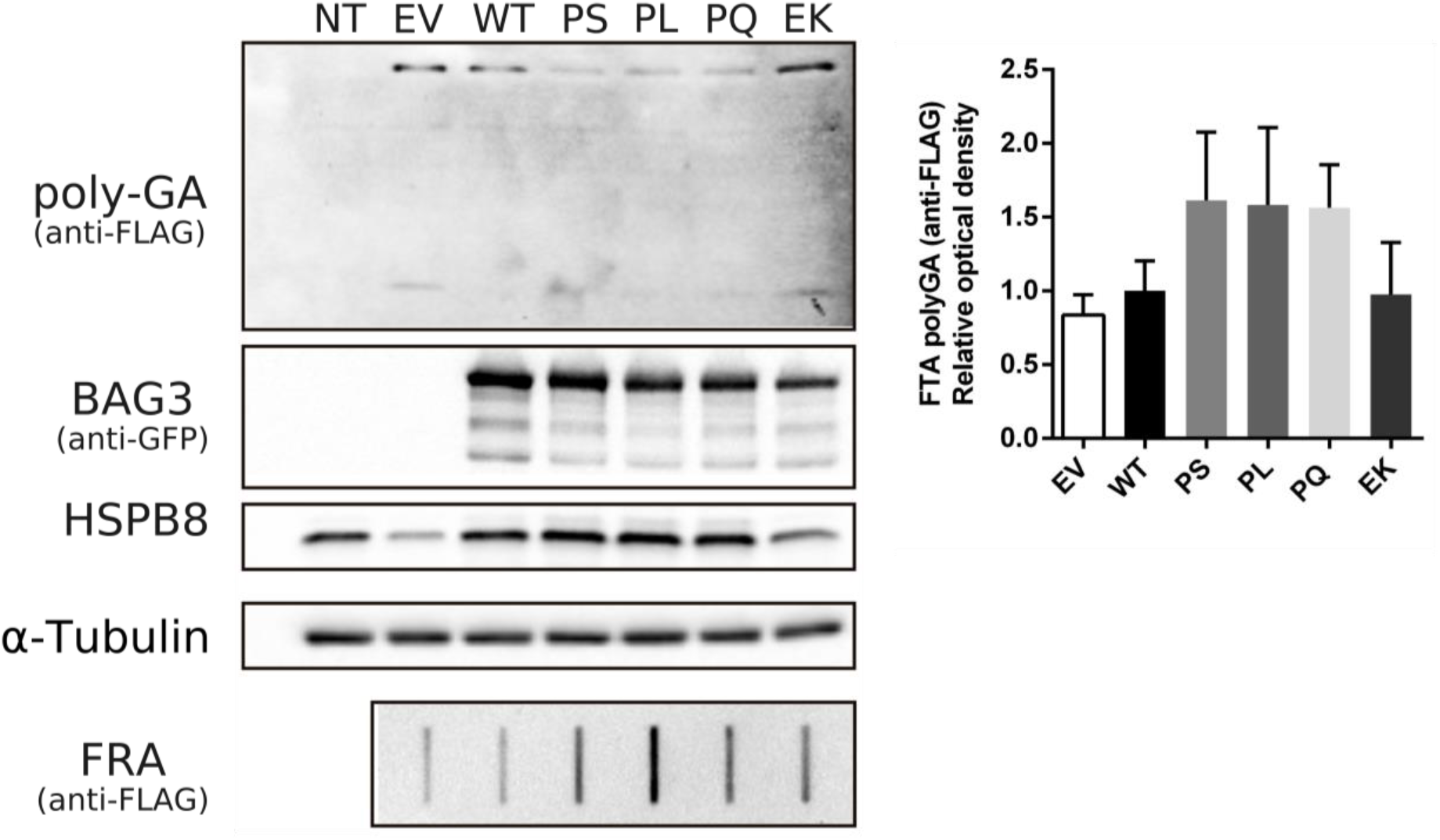
Clearance assay of neurotoxic polyGA dipeptide repeat proteins by wild type and mutant BAG3 complexes. HEK293T cells that stably overexpress HSPB8-V5 were transiently transfected with mutant BAG3-GFP constructs and FLAG-tagged polyGA. Protein lysates were collected and analysed for polyGA aggregation using a filter retardation assay (FRA). Abbreviations: non-transfected (NT), empty vector (EV), wild type (WT), Pro209Ser (PS), Pro209Leu (PL), Pro209Gln (P209Q), Glu455Lys (E455K), filter retardation assay (FRA). (n=6)

## References

1. Adriaenssens E, Geuens T, Baets J, Echaniz-Laguna A, Timmerman V (2017) Novel insights in the disease biology of mutant small heat shock proteins in neuromuscular diseases. Brain 140(10):2541–2549. https://doi.org/10.1093/brain/awx187

2. Andersen AG, Fornander F, Daa H, Krag T, Straub V, Duno M, Vissing J (2018) BAG3 myopathy is not always associated with cardiomyopathy. Neuromuscul Disord 28(9):798–801. https://doi.org/10.1016/n.nmd.2018.06.019

3. Arndt V, Dick N, Tawo R, Dreiseidler M, Wenzel D, Hesse M, Fürst DO, Saftig P, Saint R, Fleischmann BK, Hoch M, Höhfeld J (2010) Chaperone-Assisted Selective Autophagy is essential for muscle maintenance. Curr Biol 20(2), 143–148 https://doi.org/10.1016/j.cub.2009.11.022

4. Balchin D, Hayer-Hartl M, Hartl FU (2014) In vivo aspects of protein folding and quality control. Science 353(6294):aac4354. https://doi.org/10.1126/science

5. Behl C (2016) Breaking BAG: The Co-Chaperone BAG3 in Health and Disease. Trends Pharmacol Sci 37(8):672–88. https://doi.org/10.1016/j.tips.2016.04.007

6. Benoy V, Van Helleputte L, Prior R, d’Ydewalle C, Haeck W, Geens N Scheveneels W, Schevenels B, Cader ZM, Talbot K, Kozikowski AP, Vanden Berghe P, Van Damme P, Robberecht W, Van Den Bosch L (2018) HDAC6 is a therapeutic target in mutant GARS-induced Charcot-Marie-Tooth Disease. Brain 141(3):673–687. https://10.1093/brain/awx375

7. Bolognesi B, Faure AJ, Seuma M, Schmiedel JM, Tartaglia GG, Lehner B (2019) The mutational landscape of a prion-like domain. BioRxiv 1; 592121. http://biorxiv.org/content/early/2019/03/31/592121.abstract

8. Carra S, Seguin SJ, Lambert H, Landry J (2008) HspB8 chaperone activity toward poly(Q)-containing proteins depends on its association with Bag3, a stimulator of macroautophagy. J Biol Chem 283(3):1437–44. https://doi.org/10.1074/jbc.M706304200

9. Carra S, Alberti S, Benesch JLP, Boelens W, Buchner J, Carver JA, Cecconi C, Ecroyd H, Gusev N, Hightower LE, Klevit RE, Lee HO, Liberek K, Lockwood B, Poletti A, Timmerman V, Toth ME, Vierling E, Wu T, Tanguay RM (2019) Small heat shock proteins: multifaceted proteins with important implications for life. Cell Stress Chaperones 295–308. https://doi.org/10.1007/s12192-019-00979-z

10. Cha-Molstad H, Yu JE, Feng Z, Lee SH, Kim JG, Yang P, Han B, Sung KW, Yoo YD, Hwang J, McGuire T, Shim SM, Song HD, Ganipisetti S, Wang N, Jang JM, Lee MJ, Kim SJ, Lee KH, Hong JT, Ciechanover A, Mook-Jung I, Kim KP, Xie XQ, Kwon YT, Kim BY (2017) p62/SQSTM1/Sequestosome-1 is an N-recognin of the N-end rule pathway which modulates autophagosome biogenesis. Nat Commun 8(1):102. https://doi.org/10.1038/s41467-017-00085-7

11. Ciuffa R, Lamark T, Tarafder AK, Guesdon A, Rybina S, Hagen WJH, Johansen T, Sachse C (2015) The selective autophagy receptor p62 forms a flexible filamentous helical scaffold. Cell Rep 11(5):748–58. https://doi.org/10.1016/j.celrep.2015.03.062

12. Cremades N, Cohen SIA, Deas E, Abramov AY, Chen AY, Orte A, Sandal M, Clarke RW, Dunne P, Aprile FA, Bertoncini CW, Wood NW, Knowles TP, Dobson CM, Kleneman D (2012) Direct observation of the interconversion of normal and toxic forms of α-synuclein. Cell 149(5):1048–59. https://doi.org/10.1016/j.cell.2012.03.037

13. Cristofani R, Crippa V, Rusmini P, Cicardi ME, Licata NV, Sala G, Giorgetti E, Grunseich C, Galbiati M, Piccolella M, Messi E, Ferrarese C, Carra S, Poletti A (2017) Inhibition of retrograde transport modulates misfolded protein accumulation and clearance in motoneuron diseases. Autophagy 13(8):1280–1303. https://doi.org/10.1080/15548627.2017.1308985

14. Cristofani R, Crippa V, Vezzoli G, Rusmini P, Galbiati M, Cicardi ME, Meroni M, Ferrari V, Tedesco B, Piccolella M, Messi M, Carra S, Poletti A (2018) The small heat shock protein B8 (HSPB8) efficiently removes aggregating species of dipeptides produced in C9ORF72-related neurodegenerative diseases. Cell Stress Chaperones 23(1):1–12. https://doi.org/10.1007/s12192-017-0806-9

15. Crippa V, Sau D, Rusmini P, Boncoraglio A, Onesto E, Bolzoni E, Galbiati M, Fontana E, Marino M, Carra S, Bendotti C, De Biasi S, Poletti A (2010) The small heat shock protein B8 (HspB8) promotes autophagic removal of misfolded proteins involved in amyotrophic lateral sclerosis (ALS). Hum Mol Genet 19(17), 3440–3456. https://doi.org/10.1093/hmg/ddq257

16. Doong H, Price H, Kim YS, Gasbarre C, Probst J, Liotta LA, Blanchette J, Rizzo K, Kohn E (2000) CAIR-1/BAG-3 forms an EGF-regulated ternary complex with phospholipase C-γ and Hsp70/Hsc70. Oncogene 19(38):4385–95. https://doi.org/10.1038/sj.onc.1203797

17. Escusa-Toret S, Vonk W, Frydman J (2013) Spatial sequestration of misfolded proteins by a dynamic chaperone pathway enhances cellular fitness during stress. Nat Cell Biol 15(10):1–13. https://doi.org/10.1038/ncb2838

18. Fang X, Bogomolovas J, Wu T, Zhang W, Liu C, Veevers J, Stroud MJ, Zhang Z, Ma X, Mu Y, Lao DH, Dalton ND, Gu Y, Wang C, Wang M, Liang Y, Lange S, Ouyang K, Peterson KL, Evans SM, Chen J (2017) Loss-of-function mutations in co-chaperone BAG3 destabilize small HSPs and cause cardiomyopathy. J Clin Invest 127(8):3189–200. https://doi.org/10.1172/JCI94310

19. Fang X, Bogomolovas J, Zhou PS, Mu Y, Ma X, Chen Z, Zhang L, Zhu M, Veevers J, Ouyang K, Chen J (2019) P209L mutation in BAG3 dopes not cause cardiomyopathy in mice. Am J Physiol Heart Circ Physiol 316(2):H392–H399. https://doi.org/10.1152./ajpheart.00714.2018

20. Fang Y, Tsai K, Chang Y, Kao P, Woods R, Kuo PH, Wu CC, Liao JY, Chou SC, Lin V, Jin LW, Yuan HS, Cheng IH, Tu PH, Chen YR (2014) Full-length TDP-43 forms toxic amyloid oligomers that are present in frontotemporal lobar dementia-TDP patients. Nat Commun 12;5:4824. https://doi.org/10.1038/ncomms5824

21. Fernandez-Escamilla AM, Rousseau F, Schymkowitz J, Serrano L (2004) Prediction of sequence-dependent and mutational effects on the aggregation of peptides and proteins. Nat Biotechnol 22(10):1302–1306. https://doi.org/10.1038/nbt1012

22. Fischle W, Emiliani S, Hendzel MJ, Nagase T, Nomura N, Voelter W, Verdin E (1999) A new family of human histone deacetylases related to Saccharomyces cerevisiae HDA1p. J Biol Chem 274(17):11713–11720. https://doi.org/10.1074/jbc.274.17.11713

23. Fuchs M, Poirier DJ, Seguin SJ, Lambert H, Carra S, Charette SJ, Landry J (2010). Identification of the key structural motifs involved in HspB8/HspB6-Bag3 interaction. Biochem J 425(1):245–57. https://doi.org/10.1042/BJ20090907

24. Fujita K, Maeda D, Xiao Q, Srinivasula SM (2011) Nrf2-mediated induction of p62 controls Toll-like receptor-4-driven aggresome-like induced structure formation and autophagic degradation. Proc Natl Acad Sci USA 108(4):1427–1432. https://doi.org/10.1073/pnas.1014156108

25. Gamerdinger M, Kaya AM, Wolfrum U, Clement AM, Behl C (2011) BAG3 mediates chaperone-based aggresome-targeting and selective autopaghy of misfolded proteins. EMBO Rep 12(2):149–56. https://doi.org/10.1038/embor.2010.203

26. Gloge F, Becker AH, Kramer G, Bukau B (2014) Co-translational mechanisms of protein maturation. Curr Opin Struct Biol 24(1):24–33. https://doi.org/10.1016/j.sbi.2013.11.004

27. Guilbert SM, Lambert H, Rodrigue M-A, Fuchs M, Landry J, Lavoie JN (2018) HSPB8 and BAG3 cooperate to promote spatial sequestration of ubiquitinated proteins and coordinate the cellular adaptive response to proteasome insufficiency. FASEB J 0(0):fj.201700558RR. https://doi.org/10.1096/fj.201700558RR

28. Guo W, Naujock M, Fumagalli L, Vandoorne T, Baatsen P, Boon R, Ordovás L, Patel A, Welters M, Vanwelden T, Geens N, Tricot T, Benoy V, Steyaert J, Lefebvre-Omar C, Boesmans W, Jarpe M, Sterneckert J, Wegner F, Petri S, Bohl D, Vanden Berghe P, Robberecht W, Van Damme P, Verfaillie C, Van Den Bosch L (2017) HDAC6 inhibition reverses axonal transport defects in motor neurons derived from FUS-ALS patients. Nat Commun 8(1):861. https://doi.org/10.1038/s41467-017-00911-y

29. Homma S, Iwasaki M, Shelton GD, Engvall E, Reed JC, Takayama S (2006) BAG3 deficiency results in fulminant myopathy and early lethality. Am J Pathol 169(3):761–73. https://doi.org/10.2353/ajpath.2006.060250

30. Hubbert C, Guardiola A, Shao R, Kawaguchi Y, Ito A, Nixon A, Yoshida M, Wang XF, Yao TP (2002) HDAC6 is a microtubule-associated deacetylase. Nature 417(6887):455–458. https://doi.org/10.1038/417455a

31. Jiang J, Ballinger CA, Wu Y, Dai Q, Cyr DM, Paterson C (2001) CHIP is a U-box-dependent E3 ubiquitin ligase: identification of Hsc70 as a target for ubiquitylation. J Biol Chem 276(46):42938–44. https://doi.org/10.1074/jbc.M101968200

32. Johnston JA, Ward CL, Kopito RR (1998) Aggresomes: a cellular response to misfolded proteins. J Cell Biol 143(7): 1883–1898. https://doi.org/10.1083/jcb.143/7/1883

33. Kampinga HH, Craig EA (2010) The HSP70 chaperone machinery: J proteins as drivers of functional specificity. Nat Rev Mol Cell Biol 11(8):579–92. https://doi.org/10.1038/nrm2941

34. Kawaguchi Y, Kovacs JJ, McLaurin A, Vance JM, Ito A, Yao TP (2003) The deacetylase HDAC6 regulates aggresome formation and cell viability in response to misfolded protein stress. Cell 115(6):727–738. https://10.1016/s0092-8674(03)00939-5

35. Lamark T, Brech A, Outzen H, Perander M, Stenmark H, Johansen T (2005) p62/SQSTM1 forms protein aggregates degraded by autophagy and has a protective effect on huntingtin-induced cell death. J Cell Biol 171(4):603–14. https://doi.org/10.1083/jcb.200507002

36. Long M, Li X, Li L, Dodson M, Zhang DD, Zheng H (2017) Multifunctional p62 Effects Underlie Diverse Metabolic Diseases. Trends Endocrinol Metab 28(11):818–30. https://doi.org/10.1016/j.tem.2017.09.001

37. Matsuyama A, Shimazu T, Sumida Y, Saito A, Yoshimatsu Y, Osada H, Komatsu Y, Nishino N, Khochbin S (2002) In vivo destabilization of dynamic microtubules by HDAC6-mediated deacetylation. EMBO J 21(24):6820–6831. https://doi.org/10.1093/emboj/cdf682

38. McDonough H, Patterson C (2003) CHIP: a link between the chaperone and proteasome systems. Cell Stress Chaperones 8(4):303–8. https://doi.org/10.1379/1466-1268(2003)008<0303:calbtc>2.0co;2

39. Meister-Boekema M, Freilich R, Jagadeesan C, Rauch JN, Bengoechea R, Motley WW, Kuiper EEF, Minoia M, Furtado G, Van Waarde MAWH, Bird SJ, Rebelo A, Zuchner S, Pytel P, Scherer SS, Morelli FF, Carra S, Weihl CC, Bergink S, Gestwicki JE, Kampinga HH (2018) Myopathy associated BAG3 mutations lead to protein aggregation by stalling Hsp70 networks. Nat Commun 9(1):5342. https://doi.org/10.1038/s41467-018-07718-5

40. Meriin AB, Naraynan A, Meng L, Alexandrov I, Varelas X, Cissé II, Sherman MY (2018) Hsp70-Bag3 complex is a hub for proteotoxicity-induced signaling that controls protein aggregation. Proc. Natl. Acad. Sci 115(30), E7043–E7052. https://doi.org/10.1073/pnas.1803130115

41. Minoia M, Boncoraglio A, Vinet J, Morelli FF, Brunsting JF, Poletti A, Krom S, Reits E, Kampinga HH, Carra S (2014) BAG3 induces the sequestration of proteasomal clients into cytoplasmic puncta. Autopaghy 10:9, 1603–1621. https://doi.org/10.4161/auto.29409

42. Morelli FF, Mediani L, Helden L, Bertacchini J, Bigi I, Carra AD, Vinet J, Carra S (2017) An interaction study in mammalian cells demonstrates weak binding of HSPB2 to BAG3, which is regulated by HSPB3 and abrogated by HSPB8. Cell Stress Chaperones 22(4):531–40. https://doi.org/10.1007/s12192-017-0769-x

43. Murata S, Minami Y, Minami M, Chiba T, Tanaka K (2001) CHIP is a chaperone-dependent E3 ligase that ubiquitylates unfolded protein. EMBO Rep 2(12):1133–8. https://doi.org/10.1093/embo-reports/kve246

44. Prior R, Helleputte L, Van Kling YE, Van den Bosch L (2018) HDAC6 as a potential therapeutic target for peripheral nerve disorders. Expert Opin. Ther. Targets 22(12):993–1007. https://doi.org/10.1080/14728222.2018.1541235

45. Rauch JN, Zuiderweg ERP, Gestwicki JE (2016) Non-canonical Interactions between heat shock cognate protein 70 (Hsc70) and Bcl2-associated anthanogene (BAG) Co-chaperones are important for client release. J Biol Chem 291(38):19848–57. https://doi.org/10.1074/jbc.M116.742502

46. Rauch JN, Tse E, Freilich R, Mok SA, Makley LN, Southworth DR, Gestwicki JE (2017) BAG3 is a modular, scaffolding protein that physically links heat shock prtein 70 (Hsp70) to the small heat shock proteins. J Mol Biol 429(1):128–41. https://doi.org/10.1016/j.jmb.2016.11.013

47. Rusmini P, Cristofani R, Galbiati M, Cicardi ME, Meroni M, Ferrari V, Vzzoli G, Tedesco B, Messi E, Piccolella M, Carra S, Crippa V, Poletti A (2017) The role of the heat shock protein B8 (HSPB8) in motoneuron diseases. Front Mol Neurosci 10(June):1–9. https://doi.org/10.3389/fnmol.2017.00176

48. Sandell S, Huovinen S, Palmio J, Raheem O, Lindfors M, Zhao F, Haapasalo H, Udd B (2016) Diagnostically important muscle pathology in DNAJB6 mutated LGMD1D. Acta Neuropathol Commun 5;4:9. https://doi.org/10.1186/s40478-016-0276-9

49. Sarparanta J, Jonson PH, Golzio C, Sandell S, Luque H, Screen M, McDonald K, Stajich JM, Mahjneh I, Vihola A, Raheem O, Penttilä S, Lehtinen S, Huovien S, Palmio J, Tasca G, Ricci E, Hackman P, Hauser M, Katsanis N, Udd B (2012) Mutations affecting the cytoplasmic functions of the co-chaperone DNAJB6 cause limb-girdle muscular dystrophy. Nat Gen 44(4):450–S2. https://doi.org/10.1038/ng.1103

50. Schindelin J, Arganda-Carreras I, Frise E, Kaynig V, Longair M, Pietzsch T, Preibisch S, Rueden C, Saalfeld S, Schmid B, Tinevez JY, White DJ, Hartenstein V, Eliceiri K, Tomancak P, Cardona A (2012) Fiji: an open-source platform for biological-image analysis. Nat Methods 9(7):676–82. https://doi.org/10.1038/nmeth.2019

51. Schneider CA, Rasband WS, Eliceiri KW (2012) NIH Imagje to ImageJ: 25 years of image analysis. Nat Methods 9(7):671–5. https://doi.org/10.1038/nmeth.2089

52. Selcen D, Muntoni F, Burton BK, Pegoraro E, Sewry C, Bite AV, Engel AG (2009) Mutation in BAG3 causes severe dominant childhood muscular dystrophy. Ann Neurol 65(1):83–9. https://doi.org/10.1002/ana/21553

53. Semmler AL, Sacconi S, Bach JE, Liebe C, Bürmann J, Kley RA, Ferbert A, Anderheiden R, Van den Bergh P, Martin JJ, De Jonghe P, Neuen-Jacob E, Müller O, Deschauer M, Bergmann M, Schröder JM, Vorgerd M, Schulz JB, Weis J, Kress W, Claeys KG (2014) Unusual multisystemic involvement and a novel BAG3 mutation revealed by NGS screening in a large cohort of myofibrillar myopathies. Orphanet J Rare Dis 9(1). https://doi.org/10.1186/s13023-014-0121-9

54. Shy M, Rebelo AP, Feely SM, Abreu LA, Tao F, Swenson A, Bacon C, Zuchner S (2018) Mutations in BAG3 cause adult-onset Charcot-Marie-Tooth disease. J Neurol Neurosurg Psychiatry 3(89):313–5. https://doi.org/10.1136/jnnp-2017-315929

55. Sormanni P, Aprile FA, Vendruscolo M (2015) The CamSol method of rational design of protein mutants with enhanced solubility. J Mol Biol 427(2):478–490. https://doi.org/10/1016/Umb.2014/09.026

56. Takayama S, Sato T, Krajewski S, Kochel K, Irie S, Milian JA, Reed JC (1995) Cloning and functional analysis of BAG-1: a novel Bcl-2 binding protein with anti-cell death activity. Cell 80(2):279–84. https://doi.org/10.1016/0092-8674(95)90410-7

57. Takayama S, Bimston DN, Matsuzawa S-I, Freeman BC, Aime-Sempe C, Xie Z, Morimoto RI, Reed JC (1997) BAG-1 modulates the chaperone activity of Hsp70/Hsc70. EMBO J 16(16):4887–96. https://doi.org/10.1093/emboj/16.16.4887

58. Takayama S, Xie Z, Reed JC (1999) An evolutionarily conserved family of Hsp70/Hsc70 molecular chaperone regulators. J Biol Chem 274(2):781–6. https://doi.org/10.1074/jbc.274.2.781

59. Takayama S, Reed JC (2001) Molecular chaperone targeting and regulation by BAG family proteins. Nat Cell Biol 3(10):E237–41. https://doi.org/10.1038/ncb1001-e237

60. Villard E, Perret C, Gary F, Proust C, Dilanian G, Hengstenberg C, et al (2011) A genome-wide association study identifies two loci associated with heart failure due to dilated cardiomyopathy. Eur Heart J 32(9):1065–76. https://doi.org/10.1093/eurheartj/ehr105

61. Weihl CC, Udd B, Hanna M, ENMC workshop study group (2018) 234th ENMC international workshop: chaperone dysfunction in muscle disease December 8-10th 2017, Naarden Netherlands. Neuromuscul Disord 28(12):1022–1030. https://10.1016/j.nmd.2018.09.004

62. Whiten Dr, San Gil R, McAlary L, Yerbury JJ, Ecroyd H, Wilson MR (2016) Rapid flow cytometric measurement of protein inclusions and nuclear trafficking. Sci Rep 6:1–9. https://doi.org/10.1038/srep31138

63. d’Ydewalle C, Krishnan J, Chiheb DM, Van Damme P, Irobi J, Kozikowski AP, Vanden Berghe P, Timmerman V, Robberecht W, Van Den Bosch L (2011) HDAC6 inhibitors reverse axonal loss in a mouse model of mutant HSPB1-induced Charcot-Marie-Tooth disease. Nat Med 17(8):968–974. https://doi.org/10.1038/nm.2396

64. Van Helleputte L, Kater M, Cook DP, Eykens C, Rossaert E, Haeck W, Jaspers T, Geens N, Vanden Bergh P, Gysemans C, Mathie C, Robberecht W, Van Damme P, Cavaletti G, Jarpe M, Van Den Bosch L (2017) Inhibition of Histone Deacetylase 6 (HDAC6) protects against vincristine-induced peripheral neuropathies and inhibits tumor growth. Neurobiol Dis 105:300–320. https://doi.org/10.1016.j.nbd.2017.11.011

65. Youn D-Y, Lee D-H, Lim M-H, Yoon J-S, Lim JH, Jung SE, Yeum CE, Park CW, Youn HJ, Lee JS, Ikawa M, Okabe M, Tsujimoto Y, Lee JH (2008) Bis deficiency results in early lethality with metabolic deterioration and involution of spleen and thymus. AJP Endocrinol Metab 295(6):E1349–57. https://doi.org/10.1152/ajpendo.90704.2008

66. Zhu C, Beck MV, Griffith JD, Deshmuckh M, Dokholyan NV (2018) Large SOD1 aggregates, unlike trimeric SOD1, do not impact cell viability in a model of amyotrophic lateral sclerosis. 115(18):4661–5. https://doi.org/10.1073/pnas.1800187115

